# Development of the invertebrate faunas of anthropogenic habitats

**DOI:** 10.64898/2025.12.02.691838

**Authors:** Joshua M. Sammy, Jack Hatfield, Andrew Salisbury, Chris D. Thomas

## Abstract

**Aim:** We aimed to determine the relationships between species’ association with humans, their levels of habitat specialisation, and the extent to which their geographic ranges have increased or decreased.

**Location:** Great Britain.

**Time period:** Present (1981-2020)

**Major taxa studied:** Terrestrial invertebrates (n=1,722 species).

**Methods:** We determined the habitat associations for each of 1,722 species from 14 taxonomic groups in each of 18 land cover types in Great Britain. We used these values to calculate a human association index (based on whether species occupy human-modified land cover types, such as urban, suburban and coniferous plantation habitats, or occur in relatively unmodified land covers, such as several coastal habitats and marshlands) and habitat specialisation index (whether they are restricted to a few land cover types, or widely distributed across different land cover types) for each species. We then investigated the relationship between human association and habitat specialisation, as well as the relationship between human association and two metrics of range change between 1981-2000 and 2001-2020.

**Results:** Contrary to previous hypotheses, we find no evidence that species associated with human-modified environments are more likely to be habitat generalists. Furthermore, human-associated species were more likely to increase. On average, the geographic distributions of the most human-associated third of species increased by 58% over the study period, whereas the least human associated third of species declined by 7.1%.

**Main conclusions:** Humans have had increasingly large impacts on the world’s ecosystems, generating an intensity gradient of human-modification, including novel (anthropogenic) environments. Our findings show that new environments have provided opportunities for species to colonise, generating faunas which include species that have become human-associated specialists. The ongoing expansions of species in ecosystems with relatively high levels of human modification are key components of the future of biodiversity in the Anthropocene.

## 1 Introduction

The Anthropocene is marked by sweeping, human-influenced changes to the global environment, primarily through anthropogenic climate change, biological invasions, pollution, and land use change (IPBES 2019). While land use change can cause the loss of many of the original species in a given location (Newbold et al. 2015, 2018), the new environments may also provide opportunities for colonisation by other species. This is potentially similar to other forms of ecological succession, in which species colonise new environments following, for example, volcanism and natural disturbances (Chang and Turner 2019).

However, anthropogenic ecosystem types (anthromes) such as arable fields and urban environments present a number of challenges for colonising species. They may contain novel mixtures of cultivated and introduced plant species, and they are often characterised by human-generated disturbances, human structures, and chemical additions such as pesticides and pollutants. All of these differ from the conditions that were present over the preceding thousands to millions of years (Hobbs et al. 2006).

Habitat generalist species are typically regarded as more able than specialists to adjust to and occupy these human modified environments (Dawson et al. 2011; Buckley and Kingsolver 2012; MacLean and Beissinger 2017). However, environmentally distinct natural habitats, such as serpentine soils or marine cold seeps, often contain specialists that are uncommon elsewhere (Brady et al. 2005; Wang et al. 2022). Given that anthropogenic environments can also be distinct, they might acquire some specialised species too, in the same way that (chemically and structurally unusual) non-native plants may be used as habitats by uncommon native insects (Padovani et al. 2020). As anthropogenic ecosystems are both relatively new and geographically expanding, species colonising them might also be expected to be increasing.

Thus, there are the following contrasting perspectives; that (i) anthropogenic ecosystems tend to be species-poor, (ii) novel environments gain species through successional colonisation, (iii) generalists can flexibly exploit new environments, and (iv) species living in distinctive environments are often specialists. To make sense of these perspectives, it is important to understand the habitat characteristics (specialists to generalists) and status (increasing to decreasing) of species associated with ecosystems that have experienced different levels of human modification.

Here, we use data for 1,722 invertebrate species found in Britain to evaluate the relationships between species trends (frequencies of occurrence) between 1981-2000 and 2001-2020, their associations with ecosystems with different levels of human modification, and the extent to which species are specialists or generalists. We conclude that species associated with human-modified ecosystems, such as urban, suburban and coniferous plantation environments, commonly have expanding distributions, and hence that anthropogenic ecosystem types are accumulating species populations.

## 2 Methods

### Biodiversity and land cover data

We used the 2019 UK Centre for Ecology and Hydrology (UKCEH) Land Cover Map (25m rasterised land parcels, GB) (Morton 2020) as the basis for our land cover map of Great Britain. Alongside this, we used biological records data, managed and provided by a range of recording schemes (Table 1). We initially considered all of the recognised national invertebrate group recording schemes that work with the UK Biological Records Centre, and applied our data selection criteria (Supplementary Methods). These resulted in the inclusion of the taxonomic groups, and numbers of species, listed in Table 1. Data were collated for the period 1981 to 2020, except that the data for butterflies only extended to 2019 (the volume of data for butterflies outweighed those for any other taxonomic group, showing robust results regardless of the absence of 2020 data). The number of records in Table 1 excludes any records that are a duplicate of species, date and location, and excludes any records that are on a hectare that experienced land use change according to the UKCEH Land Cover Change 1990-2015 map, any records that are aggregated from hectares that are not purely of one land use type, and any records that are missing a minimum temperature, growing degree days or ratio of actual to potential evapotranspiration value value.

**Table 1:** Taxonomic groups included, relevant recording schemes, numbers of unique records, species, and data sources used in the analyses.

| **Taxonomic Group** | **Recording Scheme** | **Scheme Managed By** | **Number of records in main analysis** | **Number of species in main analysis** | **Source** |
| --- | --- | --- | --- | --- | --- |
| Bees, Wasps and Ants | Bees, Wasps and Ants Recording Society | Mike Edwards and Stuart Roberts | 129161 (88589 bees, 30409 wasps, 10163 ants) | 155 (96 bees, 56 wasps, 3 ants) | Provided directly by scheme |
| Butterflies and Moths | Butterfly Conservation | Richard Fox and Nigel Bourn | 12673985 (2877739 butterflies, 9796246 moths) | 1066 (45 butterflies, 1021 moths) | Provided directly by scheme |
| Dragonflies and Damselflies | British Dragonfly Society Recording Scheme | David Hepper | 277627 | 29 | National Biodiversity Network |
| Grasshoppers and Allies | Grasshoppers and Related Insects Recording Scheme | Peter Sutton and Björn Beckmann | 6338 | 5 | National Biodiversity Network |
| Ground beetles | Ground Beetle Recording Scheme | Chris Foster | 25651 | 32 | National Biodiversity Network |
| Hoverflies | Hoverfly Recording Scheme | Stuart Ball and Roger Morris | 321829 | 118 | Provided directly by scheme |
| Ladybirds | UK Ladybird Survey | Helen Roy and Peter Brown | 55495 | 8 | Provided directly by scheme |
| Non-marine molluscs | Conchological Society of Great Britain and Ireland | Ben Rowson | 50520 | 49 | National Biodiversity Network |
| Shieldbugs and allies | Terrestrial Heteroptera Recording Scheme | Tristan Bantock | 18159 | 10 | National Biodiversity Network |
| Soldierflies and allies | Soldierflies and Allies Recording Scheme | Martin Harvey | 23899 | 1 | Provided directly by scheme |
| Spiders, Harvestmen and Pseudoscorpions | Spider Recording Scheme | Richard Gallon | 191683 | 249 | Provided directly by scheme |

The basic unit for habitat association calculations was 1 hectare, aligned to the Ordnance Survey (OS) national grid for Great Britain (but see Species distributional changes). Hence, we filtered to remove coarser scale species records (mainly 1 and 10 km resolution records). For the purposes of analysis, finer resolution species records were coarsened to 1ha resolution on the 100m OS grid. Any records that were a duplicate of species, date and 1ha-resolution location were excluded.

The UKCEH land cover map consisted of 25m x 25m grid squares. We used the raster package in R 4.0.5 (Hijmans 2021; R Core Team 2021) to aggregate squares to produce a land cover map of 100m × 100m (1ha) resolution grid squares. We filtered to remove 1 ha squares for which >1 land cover type was present in the UKCEH 2018 land cover map, to ensure that all species records were unambiguously associated with a given land cover type. We also excluded from consideration all 1ha grid squares that experienced land use change according to the UKCEH Land Cover Change 1990-2015 map, so that changes in status represent increases or decreases within a given land cover type, rather than expansions or retractions of the land cover type itself. Finally all 1ha squares that are missing the climate variables needed for our land cover association models (below), were excluded. Following this record and cell selection procedure, we only included species that still had at least 100 presence records (see Supplementary Methods, *Record Inclusion Criteria*) in the main analysis. To test the sensitivity of the results to this inclusion criterion, we repeated the analyses for species with ≥50 presence records, and then for species with ≥200 presence records (Supp. Figs. 1, 2, 3). Taxonomic groups with fewer than 5 species (ants, and soldierflies and allies) that passed these criteria were included in the overall analysis, but were not independently included in Figures 1 and 2. Numbers of species, records and grid squares lost to these filtering steps can be found in Supp. Table 1.

**Figure 1:**
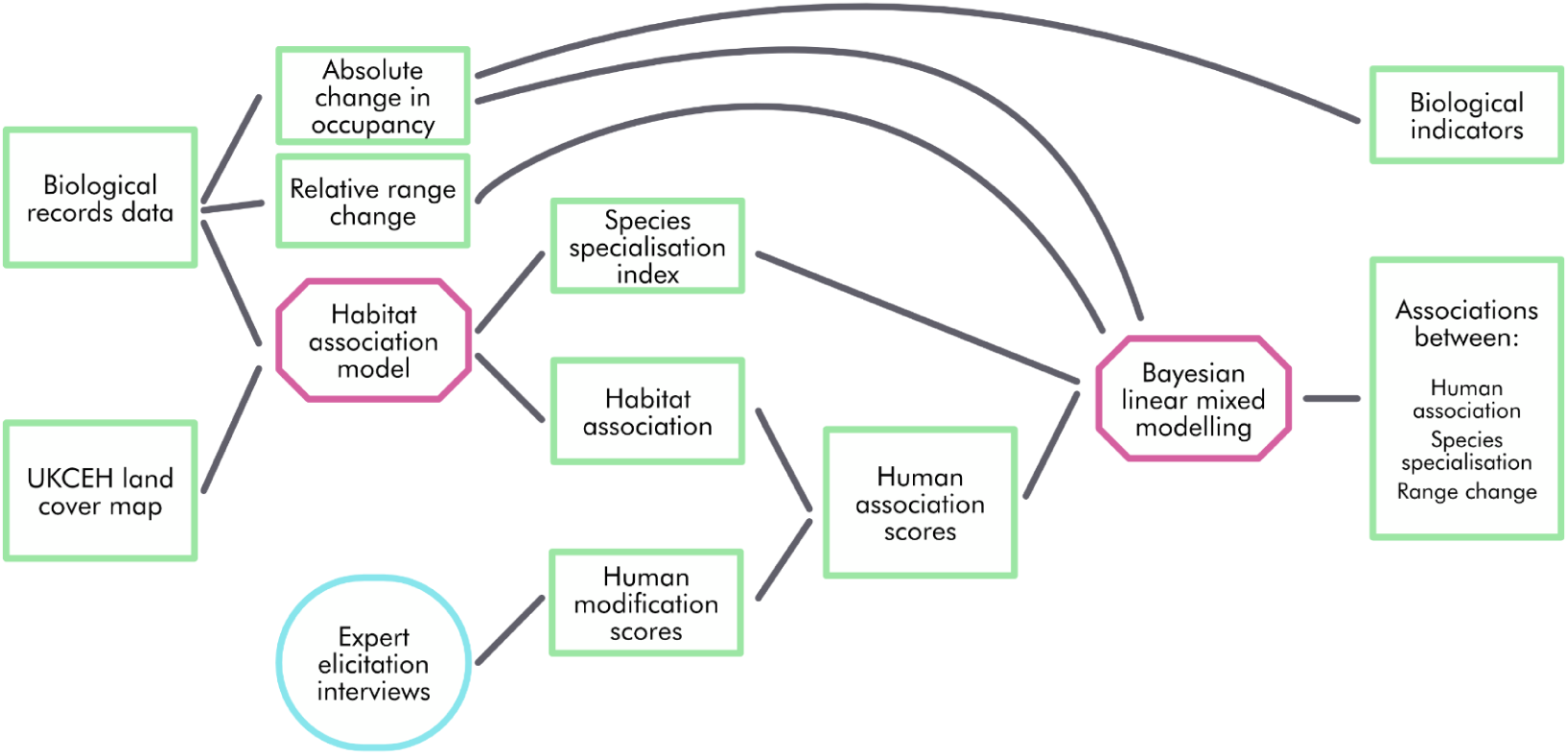
Graphical summary of paper methodology. Green rectangles represent data, pink hexagons represent models, and blue circles represent other methods.

**Figure 1:**
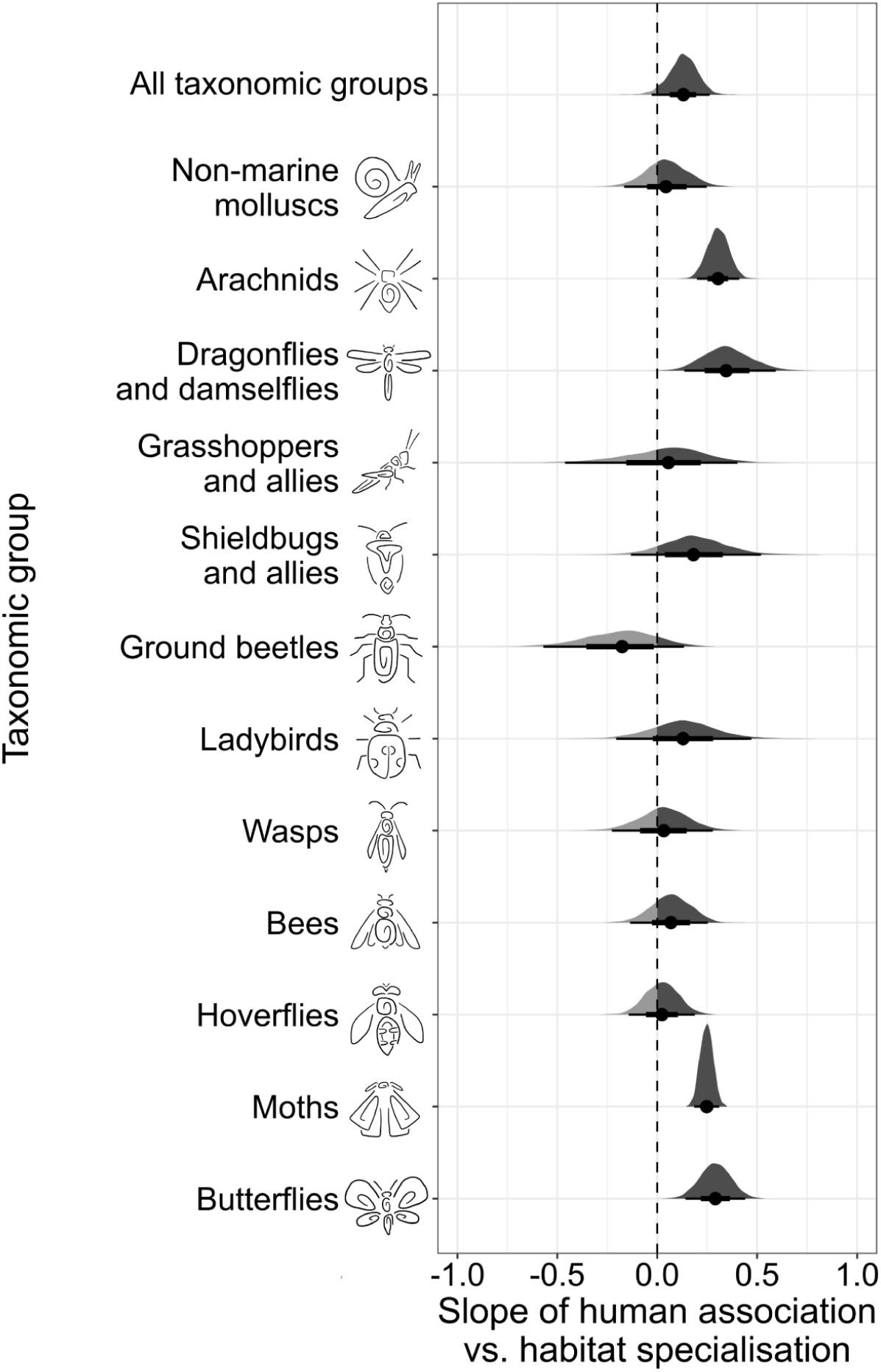
Slope of human association versus habitat specialisation, for all species pooled and by taxonomic group, from a Bayesian hierarchical model with slope and intercept as a random effect by taxonomic group. Positive slopes indicate greater specialisation in more human-modified environments. Points indicate the median estimate with the distribution of the posterior samples shown above. Error bars represent the 66% credible intervals and the 95% credible intervals. The ‘all taxonomic groups’ slope includes all groups featured in the figure, as well as ants, and soldierflies and allies, which did not contain enough species meeting data inclusion criteria (see Methods) to be included independently.

**Figure 2:**
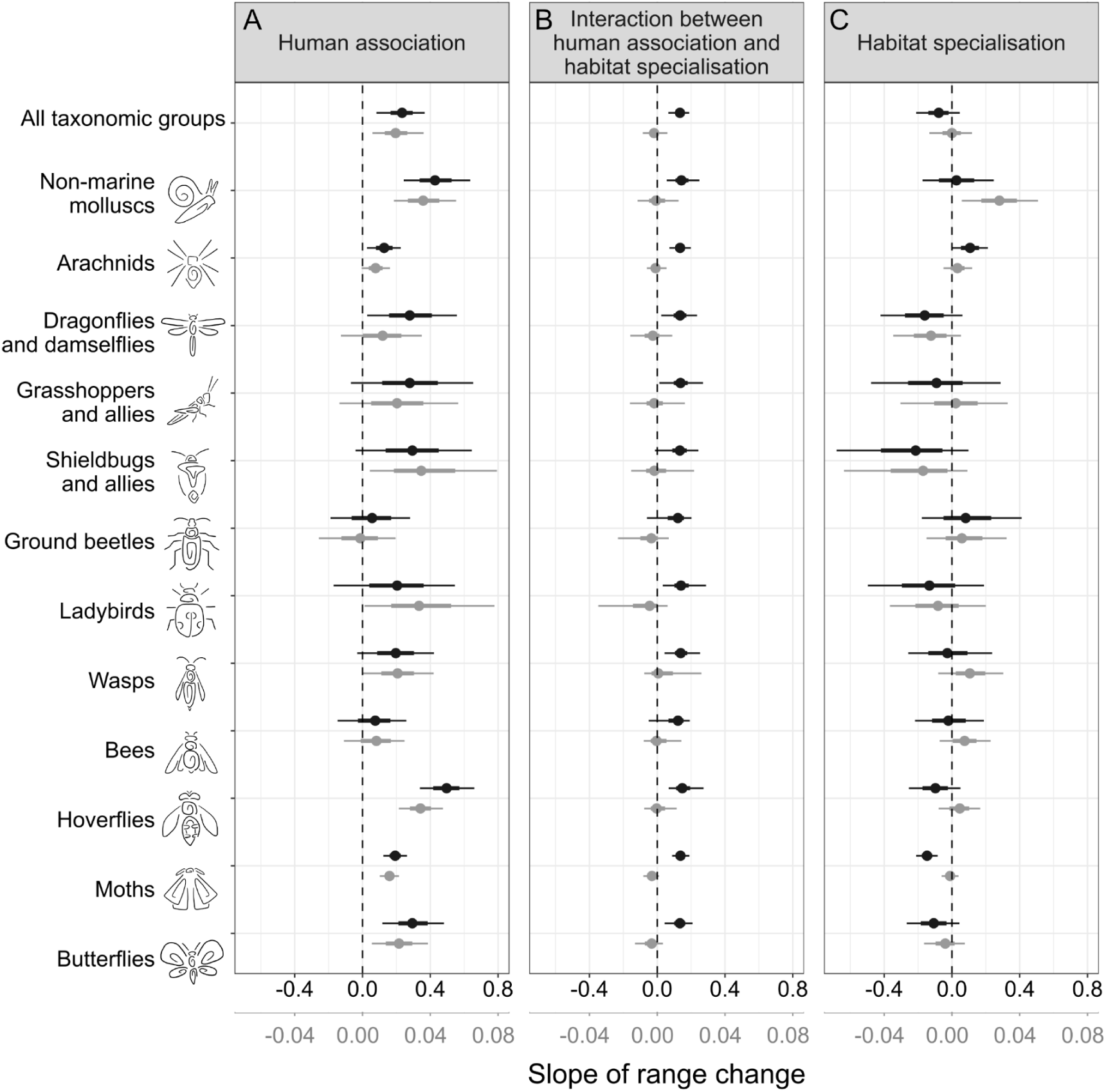
Effects of human association (A), habitat specialisation (C) and the interaction between human association and habitat specialisation (B) on range changes (slope values) by British invertebrate species between 1981-2000 and 2001-2020. Relative distribution change is in black, while change in absolute occupancy is in grey. Positive slopes in A would indicate that human associated species increased more than human avoidant species; negative slopes in C would show that habitat specialists declined most; and positive slopes in B that species specialising on the most human-modified habitats expanded most rapidly (only seen for the relative change metric). Error bars represent the 66% credible intervals and the 95% credible intervals.

### Land cover association models

We adapted the model used in Platts et al. (2019) to investigate the land cover associations and specialisation of each species (see Supplementary Methods for details). This involved the comparison of 1ha presences to pseudo-absences (pseudo-absences were presences of other species within the same taxonomic/recording scheme group) to provide an estimate of the probability that each species would be recorded in 1ha cells of each land cover type. The model includes a phenological component (only including pseudo-absences if recording events were at an appropriate time of year for each species), as well as climatic variables (minimum temperature, growing degree days or ratio of actual to potential evapotranspiration value value) so absence from particular land cover types would not be considered if those land cover types only occurred in climatically-unsuitable regions (see Supplementary Methods). Since total recording effort varies among taxonomic groups (recording schemes, Table 1), taxonomic group is included within all subsequent analyses (i.e., comparisons among species within groups are most robust).

### Species land cover specialisation index

Once land cover association probabilities were produced for each species in each land cover type, we used these data to calculate the species specialisation index. This was the coefficient of variation - the standard deviation of all of the land cover association probabilities (for a given species), divided by the mean of all of its land cover association probabilities (Julliard et al. 2006; Platts et al. 2019). Any species found disproportionately in a particular land cover would have a relatively higher value, while species found more equally in all land cover types would have a value closer to 0.

### Human modification scores

In order to quantify the levels of human modification for each land cover class, expert interviews were conducted. Five experts were selected, based on their experience with different aspects of human-environment interactions. The experts and their subject expertise are detailed in Table 2.

**Table 2:** Experts participating in the expert elicitation interviews, and their areas of expertise.

| **Expert** | **Field** | **Name** | **Areas of Expertise** |
| --- | --- | --- | --- |
| 1 | Terrestrial Ecology | Prof. Thorunn Helgason | · Microbial diversity in varying land uses<br>· Response of ecosystems to anthropogenic perturbations |
| 2 | Marine Ecology | Dr. Bryce Stewart | · Fisheries management<br>· Effects of climate change on marine ecosystems |
| 3 | Sociology | Dr. Tom O'Brien | · Environmental sociology<br>· Democratisation and its impacts on the environment |
| 4 | Historical Archaeology | Prof. Jonathan Finch | · Historic landscapes, particularly post-medieval rural landscapes<br>· Rural poverty |
| 5 | Environmental Pollution | Prof. Lisa Emberson | · The impacts of pollution on ecosystems, including agricultural ecosystems |

Experts were selected from five key fields: marine ecology, terrestrial ecology, environmental pollution, historical archaeology, and sociology. These fields were selected to represent a range of environmental interests, with historical archaeology and sociology selected to provide insight into the human element of human-environment interactions. The experts were presented with the different land cover classes used by the UKCEH land cover maps (as used in our land cover association model). These were accompanied with land cover type descriptions, based on the UKCEH land cover map documentation, most of which were derived from the UK Biodiversity Action Plan Broad Habitat definitions (Jackson 2000).

However, we adapted the land cover description slightly because our interviewees were from a range of disciplines; descriptions were also accompanied by definitions of key words and specific jargon. UKCEH descriptions were also edited to remove words or phrases that might bias the responses (for example, describing a habitat as ‘natural’ or ‘man-made’).

During the interviews, experts were asked to score each land cover class between 0 and 10, with 0 being ‘no human modification at all’ and 10 being ‘completely changed by humans’ (Supp. Fig. 4A). They were allowed to discuss their thoughts and revise their scores during the interview, and were asked to state which aspects of habitats that they considered most important when giving scores. For our analyses, we used the median of the five given human modification scores for each land cover class (Supp. Fig. 4A).

### Species human association index

We generated human association index scores for each species by calculating Spearman’s Rank Correlations between the probability of species presence in each land cover class (from land cover association models) versus the corresponding median human modification score per land cover class (as determined by the expert elicitation interviews). The rho value produced by this correlation was used as a proxy for human association in subsequent analyses (positive rho values indicating increased likelihood of presence in human-modified environments; negative human-avoidance), hereafter the ‘human association index’. Unscaled, we found generally more human associated species than human avoidant species (69% of species had positive rho values, and were human-associated; Supp. Fig. 5). This index reflects the disproportionate use of certain land cover types (equivalent to a habitat specialisation index), not other aspects of possible ecological or evolutionary specialisation (see Discussion).

### Species distributional changes

Change in species distribution was calculated as a comparison between distributions in 1981-2000 and 2001-2020. We generated two different measures of distributional change for each species, one of which was ‘relative’ (i.e., compared to other species in the same taxonomic group), and the other of which was an estimate of the ‘absolute occupancy’ of a species (i.e., likelihood of occurrence). The relative change in distribution was calculated using the methods detailed in Telfer et al. (2002) conducted in the R package sparta (August et al. 2020). The absolute change in occupancy followed the Bayesian occupancy modelling methods detailed in Outhwaite et al. (2019), and sparta (August et al. 2020) with MCMC algorithms run in JAGS (Plummer 2003).

We calculated the relative change in distribution (Telfer et al. 2002) at 10km × 10km OS grid resolution to ensure that there was sufficient data per grid square for the analysis. In contrast, the occupancy models used to estimate absolute changes in occupancy were implemented at 1km x 1km OS grid resolution as this is the scale at which they were developed and extensively tested (Outhwaite et al. 2018, 2019; Pocock et al. 2019).

Considering the different spatial resolutions of these two metrics, they differ in the extent to which they reflect more local population dynamics (at finer resolution) through to broader geographic changes to distributions (at coarser resolution). Thus, absolute occupancy modelling results at 1km x 1km resolution may emphasise population and metapopulation dynamic trends, while the relative distribution changes estimated at 10km × 10km resolution likely relates more strongly to broader-scale metapopulation and distributional trends. It is important to note that the two metrics are not directly comparable due to their difference in scale (although the results are congruent in practice; mostly linear relationship, R=0.63, R^2^ = 0.4, Supp. Fig. 6).

### Relating distribution changes, human associations and specialisation

We aimed to evaluate the relationships among the three variables: the species specialisation index, the species human association index, and distributional change (repeating the analyses for the relative and absolute measures of change). We could thereby test whether human associated species were more or less likely to be more specialised, and whether distribution changes depended on the human associations of species and their specialisation index values.

To estimate the relationship between the human association and specialisation index values of species, we constructed a Bayesian linear mixed model in Stan (Stan Development Team 2022), via the R package brms (Bürkner 2021) (Equation 1). The slope and intercept of the model were allowed to vary with the taxonomic group.

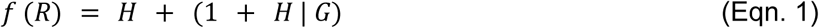

R = human association index
H = species specialisation index
G = taxonomic group

Estimating how the distribution change of species depended on their human associations and species specialisation index values using Bayesian linear mixed models, as defined in Equation 2. The modelling was repeated for both measures of distribution change: relative change and change in occupancy. We included an interaction effect between human association and species specialisation index, to determine whether range changes are experienced differently between relatively human associated and human avoidant specialists and generalists. This investigated the possibility that, for example, only human associated specialists increased.

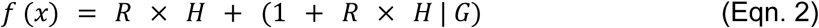

x = relative distribution change OR change in occupancy
R = human association index
H = species specialisation index
G = taxonomic group

We included measurement error in the form of standard deviation of the change in average occupancy between blocks of years for the model of change in occupancy, using the mi() function in brms (Bürkner 2017), but had no analogous error measure for relative distribution change. For each model (Equations 1, 2), we used weakly informative priors to exclude highly improbable values (Supplementary methods). The explanatory variables of human association and habitat specialisation values were scaled and centred using the scale function in R.

### Multispecies Trends

In order to visualise the changes in species through time, we constructed multi-species biodiversity indicators from the results of our occupancy models using 1,000 posterior samples from each species model, and the lambda indicator in the BRCIndicators R package (Isaac et al. 2019; August et al. 2021). This is an established biodiversity indicator method, having previously been used in reports such as the State of Nature report and UK Biodiversity Indicators (Hatfield et al. 2019; Hayhow et al. 2019; Isaac et al. 2019).

We divided species into thirds, from the most human associated third of species to the least human associated third of species, using the cut_number function from the ggplot2 package to produce three groups of equal numbers of species per group (Figure 3) (Wickham 2016). Similarly, we divided species by their associated land covers, by selecting any species that had their highest or second highest probability of presence in the land cover, to visualise the overall occupancy changes for species most strongly associated with each land cover type (Supp. Fig. 7).

**Figure 3:**
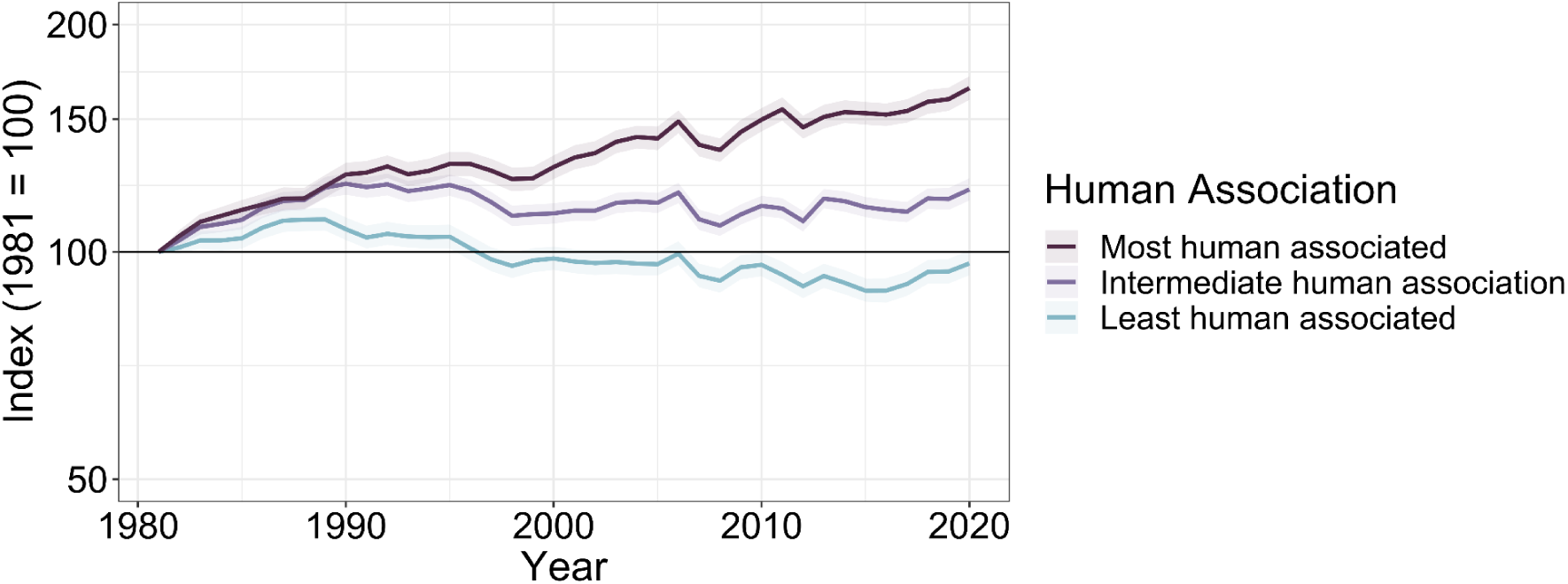
Occupancy trends for species divided into the third of species most strongly associated with human-modified environments, the middle third, and the third of species least associated with human-modified environments. Lines represent the cumulative sum of the arithmetic mean of change in log odds occupancy across species, divided into the most to least human associated thirds of species. Shading represents 95% credible intervals across the posterior samples of annual occupancy for each species.

## 3 Results

### Human Association and Habitat Specialisation

Previously, it has been hypothesised that human associated species are generalists, because generalists are the most adaptable species and thus resilient to human changes in land use (Dawson et al. 2011; Buckley and Kingsolver 2012). However, hierarchical modelling shows a high probability of a positive relationship between habitat specialisation and a species’ human association index value (95.5% of the posterior is greater than zero; Median slope = 0.13, two-tailed 95% CRI [−0.02, 0.27]) (Figure 1, Supp. Fig 8). As such, we find almost no evidence of the negative relationship which would support the hypothesis. Rather, the generalists in these locations share their environments with a suite of specialists.

This tendency for species with stronger human associations to be more specialised (positive median slope estimates) holds for 13 of the 14 taxonomic groups separately, but with varying levels of likelihood (Figure 1, Supp. Fig. 8, Supp. Table 5). This result resonates with the observation that rare insect species are sometimes associated with distinct non-native plant species (Padovani et al. 2020). Ground beetles are the only taxonomic group for which human associated species tend to be generalists (negative median slope estimate, Figure 2.1).

It is, however, important to note the variation between individual species as seen in Supplementary Figure 8. The trend for human associates to be land cover specialists does not preclude the existence of human avoidant specialists, nor of relatively human associated generalists. Rather, we show that the most human-modified environments contain *at least as* high a proportion of specialists as less modified land covers, and clearly more for arachnids, odonates (dragonflies and damselflies) and Lepidoptera (both moths and butterflies).

Our overall conclusion is robust to the exclusion of particular taxonomic groups (leaving each group out in turn; Supp. Fig. 9). It is also robust when varying the species inclusion criteria (i.e., adopting a minimum threshold of at least 50, 100 (main text) or 200 records for a species to be included in the analysis), when aggregating land cover types into broader categories, and when considering the human modification scores of different experts separately (Supp. Fig. 1).

### Species Distribution Trends

Our results reveal that the invertebrate species associated with the most human-modified environments were more likely to increase in distribution than human-avoidant species for both metrics of range change between 1981-2000 and 2001-2020 (Figure 2A, Supp. Fig. 10, Supp. Table 6).

For *relative* distribution changes, we found a one-tailed 99% probability of a positive slope for the relationship between human association and distribution change (Median slope = 0.23, two-tailed 95% CRI [0.08, 0.36]). For *absolute* occupancy changes, we also found a one-tailed 99% probability of a positive slope for the relationship between human association and change in occupancy (Median slope = 0.02, two-tailed 95% CRI [0.006, 0.4]). Thus, the conclusion that human-associated species expanded the most was robust to which of the two measures of distribution change was adopted. The ‘all taxonomic groups’ result (top row, Figure 2) is robust to the exclusion of each individual taxonomic group in turn (Supp. Fig. 11). Effect sizes are shown in relation to the indicator analysis (Figure 3, below).

Effect sizes are harder to articulate due to the derived nature of the metrics used in this analysis. Metrics such as the human association index and index of relative distribution change are useful in exploring where species lie on the spectrum in comparison to other species. As such, average effect sizes are more easily understood via the indicator analysis below (Figure 3), or through the predicted trendlines as presented in Supplementary Figures 8 and 10. We have also included a presentation of effect sizes in Supplementary Table 5.

In contrast, the results for habitat specialisation are inconsistent between the two measures of change (Figure 2C). The analyses of *absolute* change in occupancy show no effect of specialisation (slope estimates close to zero). Analysis of *relative* distribution change also shows no clear effect (Median slope = −0.07, 95% CRI [−0.20, 0.04]).

The positive interaction effect for relative distribution change indicates that specialists of human-modified ecosystems are doing better, and specialists of relatively natural environments are declining the most (Median interaction slope = 0.13, 95% CRI [0.06, 0.19], positive slopes in Figure 2B, Supp. Fig. 10).

This interaction effect was not seen in absolute change in occupancy, and as such we recommend caution in interpretation of the interaction results. Overall, the results suggest positive species distribution changes, on average, for human-associated species, but limited or mixed effects of specialisation (Supp. Table 2).

### Aggregated Assemblage Trends

Multi-species indicator methods (Hayhow et al. 2019; Isaac et al. 2019) applied to our absolute occupancy estimates (1 km x 1 km resolution) also show an average positive occurrence trajectory for human-associated species and a slight negative trend for human-avoidant species (Figure 3). Taking the average of the last five years of indicator values, we estimate that the most human-associated third of species increased on average by 58% (indicator rising to a mean of 158 in 2016-2020 compared to the base year of 1981) over the study period, whereas the least human associated declined by 7.1% (indicator declining to a mean of 92.9 in 2016-2020). When species are divided into groups according to the environments they are most strongly associated with, we see similar trends; higher occurrence trajectories for the most highly human associated environments, and lower occurrence trajectories for the least human associated environments (Supp. Fig. 7).

## 4 Discussion

Our primary conclusion is that, on average, human-associated species have been expanding in relative and absolute range sizes (frequency of occurrence) over a forty year period, while human-avoidant species have, on average, been declining. The positive association between species’ levels of habitat specialisation and their human-associations (Figure 1), and lack of an overall effect of specialisation on changes through time, provides no evidence for the frequently-held perspective that human-associates are predominantly generalists (Clavel et al. 2011; Dawson et al. 2011; Buckley and Kingsolver 2012; MacLean and Beissinger 2017; Minter et al. 2020). The empirical results suggest the reverse. Rather than being species-poor and assemblages of generalists (Clavel et al. 2011; Newbold et al. 2018), our results indicate the emergence of sets of species associated with new Anthropocene environments. They represent a suite of human associated species, including specialists, some expanding in these environments, others surviving in them (Strauss and Biedermann 2006; Ellis and Ramankutty 2008; Baldock et al. 2015; Ellis et al. 2020; Fox et al. 2022).

Of note, this analysis focuses on locations which were regarded as having unchanged land cover during the study period (locations with changing land cover were excluded, see Supplementary Methods). Hence, the increased occupancy (distributions) of human-associated species represent expansions into human-modified locations that already existed, rather than simply a reflection of certain land covers increasing over the study period. Nonetheless, habitat availability is known to facilitate range expansions (Platts et al. 2019), and land cover types with high human-modification scores do tend to be more frequent than less modified ecosystem types in Britain (Supp. Fig. 12). It is also the case that the land area of some of these anthropogenic environment types is increasing (Rounsevell and Reay 2009), so these species should additionally increase in range as more land becomes available to them.

An additional factor to consider is the potential role of increasing recording effort for some taxonomic groups in human modified land cover types (where people live). The relative recording effort in suburban environments increased between the two time periods for some taxonomic groups (wasps no change, to hoverflies with a six-fold increase), which could potentially generate positive human association-range change relationships in those groups experiencing the greatest such increases (Supp. Fig. 13). However, the level of recording increase in more or less human modified environments was unrelated to the slope of the human association/range change relationship in general, and in the specific case of increased suburban recording (Supp. Fig. 14). The positive distributional trends of human associates also remained consistent when considering varying the species inclusion criteria (i.e. 50, 100 or 200 records for species inclusion), when using species with a high number of records in every land cover class, when aggregating land cover types, and when considering the human modification scores of different experts separately (Supp. Fig. 2).

It is important to recognise that human associated specialists vary widely in their origins and present-day affinities. For example, the cellar spider (*Pholcus phalangioides*) likely originated as a subtropical cave spider that now finds adequate substitutes within thermally buffered and darkened buildings (Schäfer et al. 2001; Huber 2011), and the leopard slug *(Limax maximus*) is so widely found close to human habitations that its original distribution (presumed European) and habitat are unknown. Both species are considered to be anthropophilic (Boycott 1934; Jackson et al. 1990), and are specialists of the environments they inhabit. Other species include the native apple leaf skeletonizer moth (*Choreutis pariana*), a Eurasian species which feeds on fruit trees, and the western conifer seed bug (*Leptoglossus occidentalis*), an introduced North American species that feeds primarily on introduced conifers. These various individual requirements then map on one or more of the eighteen land cover types, which are internally heterogeneous. Hence, only a subset of 1 ha cells within a given land cover will meet the specific habitat needs of each species at any given time (for example, the larval host plants of particular Lepidoptera species do not occur everywhere within a particular land cover type).

It is also useful to distinguish between habitat and physiological specialists, although the two may be linked. For example, the cellar spider likely has thermal physiological limits that confine it to buildings, while the seed bug may be constrained by its dietary physiology to live mainly in conifer plantations and sub/urban plantings. However, other habitat specialists may be confined to one or a few land cover types as a result of avoiding natural enemies or competitors. Further research will be required to elucidate the multiple causes of specialisation in the many different kinds of human-modified environments. Nonetheless, the resulting habitat associations are important determinants of the distributions of species and, for practical policy and landscape management, awareness of human associated habitat specialists is valuable.

The existence of human associated specialists is already recognised, in the sense that many ‘specialist’ species in Britain are found disproportionately in areas with traditional farming practices and land cover types - although some have declined again following the recent intensification of agriculture in Britain (Whittingham et al. 2009; Smith et al. 2020). Presumably, such species colonised ‘earlier’ anthropogenic environments in previous centuries and millennia. The main difference between these and more recently expanding human associates of anthropogenic environments seems to be in the timing of their expansions, and the increasingly global scale of colonisation. The human associated species documented in our results are, on average, exhibiting more positive distributional trajectories, as they colonise environments that have come into existence or that have become more extensive in recent decades and centuries. The emergence and dynamics of these new biotas may help us to understand how former human-caused changes to ecosystems (including historic farming and forestry) stimulated past distributional and community changes.

These dynamics present interesting challenges for conservation. At a time of ever-increasing human influences on planetary processes and accelerating rates of distributional and community changes (Chen et al. 2011), change is the norm (Thomas 2017). As a result, conservation management inevitably becomes about maintaining and managing successive waves of human associated species, at least in the more heavily modified parts of the planet surface. The focus of much conservation management in Britain over the past 50 years has been to replicate various traditional land management practices, aiming to maintain or restore the habitats of historic human associates (Fuller et al. 1995). In contrast, encouraging the expansion of more recent arrivals is rarely considered. We suggest that species which have expanded into human modified ecosystems in recent times are no more or less ‘worthy’ of investigation and conservation than those which arrived centuries or millennia ago. Understanding the ecological requirements of the most recent wave of anthropic species and putting in place measures that accelerate their expansion may be an effective way to maintain and increase diversity levels, as an addition to protecting our historic associates.

In conclusion, contrary to expectations, species associated with the most human-modified land cover types are often relative habitat (land cover) specialists that are less well represented in ‘more natural’ ecosystem types. Furthermore, human-associated species are disproportionately increasing, providing a degree of exception with the prevailing narrative that biodiversity is declining (nearly) everywhere. Increasingly, human associated species represent the biological norm of the Anthropocene, particularly in environments that most people experience most of the time.

## Data Availability Statement

Biological Records data were provided by the various recording schemes acknowledged in Table 1. To access data, these schemes may be contacted directly through the Biological Records Centre (https://www.brc.ac.uk/recording-schemes). Data produced through the analyses conducted in this paper are available in the figshare repository. Code used to conduct the analysis in this paper is also available in the figshare repository.

## Ethical Compliance Statement

This study complies with the ethical standards, approved by the University of York Department of Biology Ethics Committee. All participants in the expert elicitation protocol provided informed consent, including for the use of their names and descriptions in this study.

## Author Contribution

All authors contributed extensively to this paper. J.S., A.S. and C.T. conceived the research, A.S. and C.T. supervised the project. J.S. conducted the analyses and wrote the paper. J.H. and J.S. conducted the Bayesian analyses. All authors discussed results, analyses and edited the paper at all stages.

## Acknowledgements

We are pleased to acknowledge the recording schemes detailed in Table 1, and their respective scheme organisers, for allowing us to access their data, the hard work they do managing these valuable datasets, and for providing advice on how species should be treated in our analysis. We are especially grateful to all of the individual volunteer recorders contributing records to each scheme.

We are grateful to Professor Helen Roy for her extensive support through all stages of the work in this project. We are grateful to Dr. Anna Woodhead for her support with the expert elicitation protocol, and the experts detailed in Table 3 for providing their expertise as part of this process. We would also like to thank Dr. Philip Platts for his support and advice through the use of the habitat association modelling protocol, and Dr. Rob Boyd for his advice on accounting for bias in the occupancy modelling.

This project analysis was undertaken on the Viking Cluster, which is a high performance compute facility provided by the University of York. We are grateful for computational support from the University of York High Performance Computing service, Viking and the Research Computing team.

We are also pleased to acknowledge the Natural England Research Council for their funding during this project, and the Leverhulme Trust Research Centre for their support through the Leverhulme Centre for Anthropocene Biodiversity.

For the purpose of open access a Creative Commons Attribution (CC BY) licence is applied to any Author Accepted Manuscript version arising from this submission.

## Supplementary

### Supplementary Methods

#### Record inclusion criteria

For this analysis, (i) we removed any grid squares where more than one land cover class was present (because 1ha species records from these cells could not be attributed unambiguously to a single land cover type). (ii) Similarly, we removed grid squares where any of the constituent 25m x 25m squares experienced a change in land cover, according to the UKCEH Land Cover Change 1990-2015 map (Rowland et al. 2020). This was to ensure that species records were associated with the correct land cover type (species records from intermediate dates again cannot be assigned unambiguously to one land cover type if the land cover of a cell changes). Thus, changes in frequencies of species reflect increases or decreases in their occurrence within a given land cover type, rather than an expansion or contraction of the areal coverage of those land cover types. (iii) We also removed any records found on littoral rock, littoral sediment, and saltwater, as littoral habitats are found between the high and low water mark, and saltwater is found below the low water mark. We considered it unlikely that any stray records (of the taxa considered) would be found as breeding populations in these locations. (iv) Finally, for the purposes of habitat association modelling, we aggregated duplicate records for each grid square so that presence was only counted once per species per grid square across the 40-year period of study (with climatic variables averaged across each of those record instances, see *Land cover association models* below).

#### Land cover association models

We compared the 1ha resolution presence records for a focal species with 1ha resolution pseudo-absences, which were presence records of other species in the same taxonomic group (but where the focal species was not observed), within a 50km radius of the presence records (to account for the effects of geographic bias). Each presence and pseudo-absence record was associated with a land cover class from the UKCEH 2018 land cover map, at 1ha resolution, as well as climatic variables (for the 1km^2^ within which the 1ha species record was located). The chosen climatic variables were the minimum temperature (Met Office et al. 2021), growing degree days above or equal to 5 degrees Celsius (Martinez-de la Torre et al. 2018) (GDD5), and the ratio of actual to potential evapotranspiration (Robinson et al. 2017; Martinez-de la Torre et al. 2018) (APET). Monthly actual evapotranspiration (AET) was downloaded from the NERC Environmental Information Data Centre (Martinez-de la Torre et al. 2018), updated to 2017 by the UK Centre for Ecology and Hydrology. Daily values for potential evapotranspiration (PET) is available publicly through the CHESS dataset (Robinson et al. 2020*a*, 2020*b*), and was averaged to month for us by the UKCEH. We then calculated APET by dividing AET by PET. GDD5 was provided to us directly by the UKCEH, derived from the CHESS dataset and averaged to month. Minimum temperature per month was accessed via the Centre for Environmental Data Analysis (CEDA) Archive (Met Office et al. 2021), and the mean of this across the year was calculated. All of these were specific to 1km^2^, and so were associated with records that fell within that area; we used the mean value for the previous five years (including the year of the record). The exception to this was for GDD5 and APET for species records after 2017. Data were not available for 2018-2020, so the five year average from 2013-2017 was instead used for these records. If a species was recorded as present in a particular 1ha grid square in multiple years, the five-year mean climate was derived for each record and the average was then taken across records. We applied the same criteria for selecting environmental variables associated with pseudo-absences.

We compared species presence records to other records of species (our pseudo-absences) within the same recording-scheme group in order to reduce the chance of false-absences, with the assumption that recorders of a particular taxonomic group are more likely to record other species in the same taxonomic group. Discussions with biological recording schemes allowed us to decide whether the taxon should be defined as the scheme itself (as was the case with arachnids), or whether it should be divided into taxonomic subgroups. For the most part, species were categorised into taxonomic groups based solely on the recording scheme from which the record came, as recorders in each scheme were considered likely to record other species within the same scheme, regardless of how closely related they were. However, bees, ants, wasps (because different experts often specialise in recording these taxa), butterflies and moths (because diurnal and nocturnal recording methods differ, and only a subset of butterfly recorders identify moths) were divided into distinct taxonomic groups.

We also reduced the number of false absences by constructing phenological models. We modelled the phenology of each species, accounting for both univoltine and multivoltine taxa to establish their seasonal activity periods. We then removed background records within the spatial buffer that were in the tails of this phenology (the lower and upper 10th quantile) (Platts et al. 2019). Thus, we did not use records of ‘other species’ as pseudo-absences if they were outside the time of year when a given focal species was in a life stage that recorders monitored.

A general linear model for each species was then constructed, with all presence/pseudo-absence records for the species, within the background buffer, against land cover class, minimum temperature, GDD5 and APET (Supp. Eqn. 1). Once this model was constructed, probabilities of presence were predicted for each land cover class, controlled for the effects of climate using the record’s associated climatic variables.

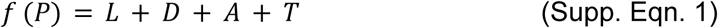

P = Presence/Absence
L = Land cover class (factor with 18 categories)
D = Growing Degree Days above 5 degrees celsius (GDD5)
A = Ratio of potential to actual evapotranspiration (APET)
T = Mean monthly minimum temperature across the past 5 years

We repeated this analysis using 9 of the 10 UKCEH aggregate land cover classes (saltwater and littoral habitats were still excluded), in place of the 18 land cover classes used in the main. For each record, we thus converted the land cover class to its corresponding aggregate land cover class (Supp. Table 2). This was to determine whether our results were strongly affected by the aggregation of land covers used. The results remained qualitatively similar, with a decrease in the positive relationship between human association and habitat specialisation (Supp. Figs. 1, 2).

#### Human modification scores

We conducted expert elicitation interviews to determine scores of human modification for each land cover class. The definition of modification was left up to each expert’s opinion, to ensure that their scoring would reflect their own understanding and field of expertise. When considering how modified an environment is, experts frequently cited accessibility of the environments to human populations in Great Britain, the history of the land cover type and recency of any change, and introduced (accidental or cultivated) species. They were also asked to state how comfortable they felt with the score they gave, this representing their knowledge of the subject, or their comfort with narrowing the habitat score to a single value (Supp. Fig. 4B). Experts were most comfortable in the scoring of arable, coniferous woodland, suburban and urban habitats, the four habitats with the highest human modification scores, and were least comfortable in the scoring of calcareous grassland and acid grassland.

There were issues with applying a human modification score to each land cover class. Notably, the land cover classes as defined by the UKCEH are likely to each cover a range of levels of human modification. Experts were asked to give a score based on what they would expect the modal grid square of the land cover class to be in the island of Great Britain over the past 40 years. The median of all scores given by all experts was 7, suggesting that they considered most environments in Great Britain to be fairly human modified. Indeed, no land cover was given a score below 2 by any expert.

The median human modification scores for each land cover class was then produced from these interviews. This was used for all of the main analyses (Supp. Fig. 4A). We compare the scores used for our study to other, established and global, human modification metrics (Supp. Fig. 15) (Ellis and Ramankutty 2008; Newbold et al. 2015; Ellis et al. 2020). We also repeated all analyses, using each expert’s human modification score for each land cover in turn to calculate species’ human association index (see *Relating distribution changes, human associations and specialisation* below). The overall results were very similar to the results using median scores (Supp. Figs. 1, 2).

When performing the repeat analysis for aggregate land covers (Supp. Table 2; Supp. Figs. 1, 2), we calculated the weighted mean human modification score for each aggregate habitat based on the number of hectares each land cover class contributed to their aggregate total.

#### Species distributional changes

We investigated the relative change in distribution following the method described in Telfer et al. (2002) using the sparta R package (August et al. 2020). For this method, we calculated changes by comparing species within the same taxonomic group, to account for variation in recording frequency between taxonomic groups. Records were used at a hectad (10km × 10km square) spatial resolution for this analysis. Species’ recording frequency was found by calculating the proportion of cells that were recorded for that taxonomic group in each time period (1981-2000 and 2001-2020) for each species. Each cell was counted only once per time period. The standardised residuals of each species from the taxonomic group’s overall change in recording frequency were then used to determine how much each species changed in range relative to others in the same taxonomic group.

The absolute change in occupancy was calculated using the occupancy modelling methods detailed in Outhwaite et al. (2019), and the sparta package (August et al. 2020) with Markov Chain Monte Carlo algorithms run in JAGS (Plummer 2003). Records in this analysis were conducted at a finer grain, monads (1km x 1km squares). For these models, prior choice was based on previous evaluation (Outhwaite et al. 2018). As in the Outhwaite et al. (2019) study we used the random walk model, half-cauchy hyperpriors for random effects, and considered list length a categorical variable (Outhwaite et al. 2019). These methods estimated the occupancy for each species based on its detection history. This was constructed using visits recording that taxonomic group within each monad. This produced a yearly estimate of the proportion of monads (with at least two records for the taxonomic group in separate years over the whole time span) that the species was present in. We then took 1,000 samples from the posterior distribution of occupancy estimates per year, calculated the mean occupancy per year, and then found the mean value of occupancy across years for 1981-2000 and 2001-2020. The change in occupancy was calculated as the difference between the mean occupancy in the latter period and the former period. For most taxonomic groups, we ran the model for 40,000 iterations with 20,000 of these as burnin and a thinning rate of three and three chains. For butterflies and moths, taxonomic groups with particularly large quantities of records, we ran the model for 20,000 iterations with 10,000 of these as burnin. Duplicate records were used if they occurred in the same location on different days, but not if there were multiple records on the same day. We assessed our occupancy models using the precision criteria available in the dataMetrics() function in sparta (August et al. 2020). The data requirements to produce precise trends from this occupancy model formulation have been rigorously tested using expert elicitation and classification trees. We used the criteria in Figure 3b of Pocock et al. (2019) and found that all of our species were compliant. As the criteria were met we can be 98% confident that the resulting trends meet the expert determined precision criteria (Pocock et al. 2019).

#### Relating distribution changes, human associations and specialisation

For each model used in calculating the relationships between human association, distribution change (relative and absolute) and habitat specialisation (Equations 2, 3), we used weakly informative priors to exclude highly improbable values.

When calculating the relationship between range change, human association and specialisation, slope and intercept were both set to a normal distribution with a mean of 0 and a standard deviation of 0.25, while standard deviation was set to an exponential with rate of 1.

When calculating the relationship between human association and specialisation, slope and intercept priors were both set to a normal distribution with a mean of 0 and a standard deviation of 1.5, while standard deviation was set to an exponential with rate of 1. Human association and habitat specialisation values were standardized to have a mean of 0 and standard deviation of 1, while relative distribution change and change in occupancy were left unscaled to allow easier derivation of effect size.

In order to investigate the influences of recording bias on our results, we calculated how much the distribution of records per taxonomic group differs from the actual distribution of land cover types. We calculated the proportion of records associated with each land cover type (for a given taxonomic group) and compared this to the proportion of hectares occupied by each land cover type in Britain overall (with littoral rock, littoral sediment and saltwater excluded from both totals). We then calculated the difference between the actual and recorded distributions of land cover type for time periods 1 and 2, negative values representing ‘under-recording’ (relative to total land cover) and positive values ‘over-recording’ (Supp. Fig. 13).

To estimate changes in recording bias, we took the difference in recording bias for each land cover (for each taxonomic group) between the two time periods (which mainly showed that recording was more biassed towards suburban land cover in the second than in the first period). We estimated the overall shift in recording towards anthropogenic habitats for each taxonomic group by calculating a Spearman’s correlation between these differences in relative recording per land cover type and the human modification score of each land cover class. A strongly positive rho is interpreted as more recording of that taxonomic group in more anthropogenic habitats in the second period; a negative correlation as increased recording in relatively natural environments in the second time period.

If this change was generating the observed results, then we predicted that taxonomic groups with the greatest increases (between periods 1 and 2) in recording bias towards (i) suburban habitats, and (ii) towards anthropogenic land cover types in general would show the strongest positive links between species human associations and their distribution changes (the slope values from Figure 2). There was no such tendency, whether considering a taxonomic group’s bias towards suburban environments (Supp. Fig. 14A) or towards human environments in general (Supp. Fig. 14B).

#### Multispecies Trends

When calculating multispecies trends, the start year is fixed to a value of 100 (as is convention with biodiversity indicators). Subsequent values are calculated using the arithmetic mean of the change in log odds (Hatfield et al. 2019; Isaac et al. 2019). If a species was not detected for more than 5 years at the start of the time series then only estimates after the first detection were used, predominantly to account for introduced species.

### Supplementary Figures and Tables

**Supplementary Figure 1:**
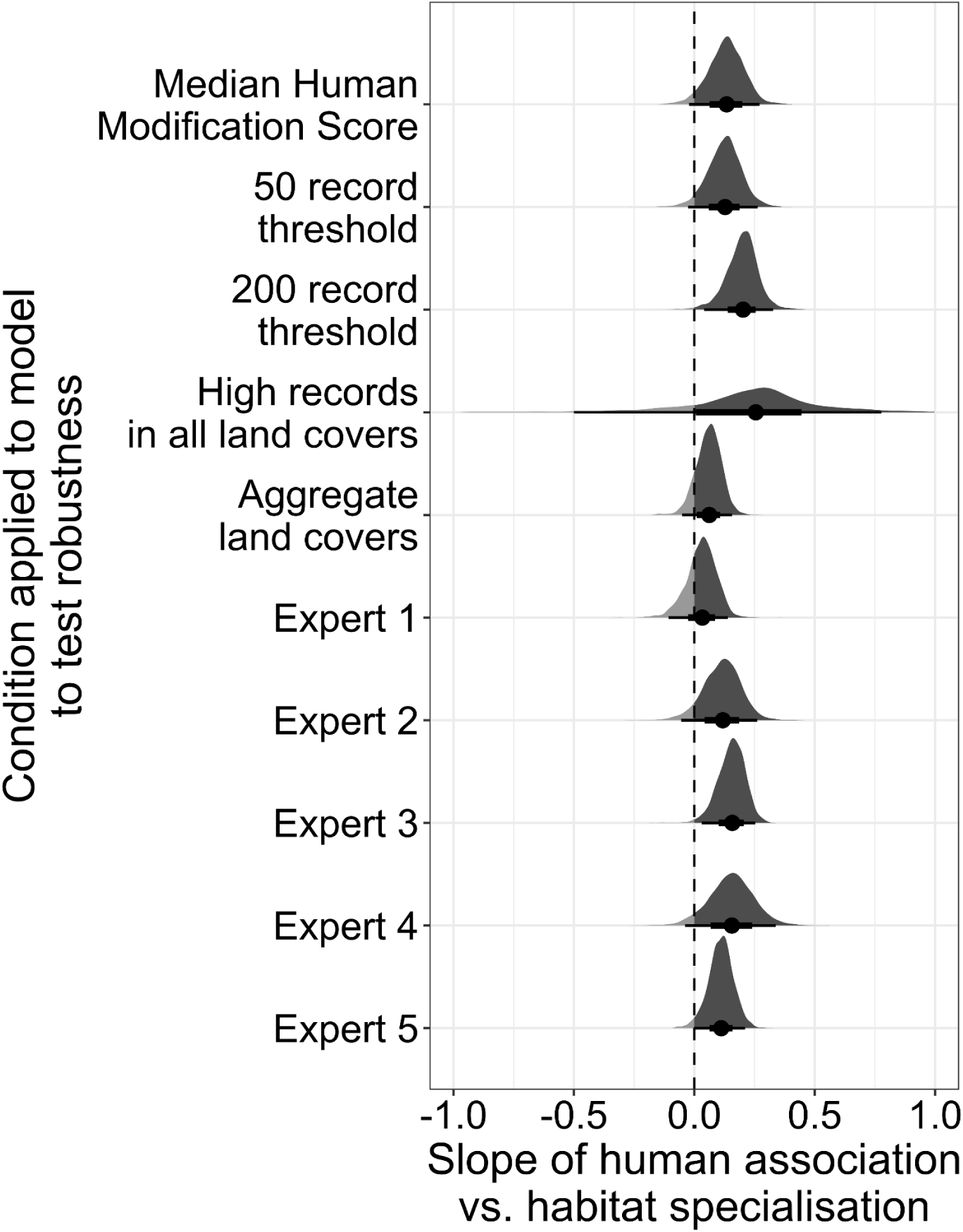
Slope of human association versus habitat specialisation, for all species pooled, from a Bayesian hierarchical model with slope and intercept as a random effect by taxonomic group. comparing the effects of various tests for robustness. These include: 1) Adjusting the thresholds for species selection to species with 50 presence records and 200 presence records (as compared to the 100 presence record threshold used in the main; top row median, here), 2) Adjusting the threshold for species selection to species with 15 records (presence or absence) in each of the 18 land cover classes, 3) Aggregating land cover classes according to the UKCEH Aggregate Land Cover Classes (Morton 2020) (Supp. Table 2), 4) Calculating human association index based solely on the human modification scores of each expert in turn (see Methods). Positive slopes indicate greater specialisation in more human-modified environments. Error bars represent the 66% credible interval and the 95% credible interval. There is some effect of condition on slope, with a modest shift towards a more positive relationship with a threshold of 200 presence records per species, and a shift towards a more negative relationship using Expert 1’s habitat modification scores or the aggregate land cover classes. However, these do not qualitatively change our overall conclusion, that human associated species are not primarily generalists, giving us confidence in our findings.

**Supplementary Figure 2:**
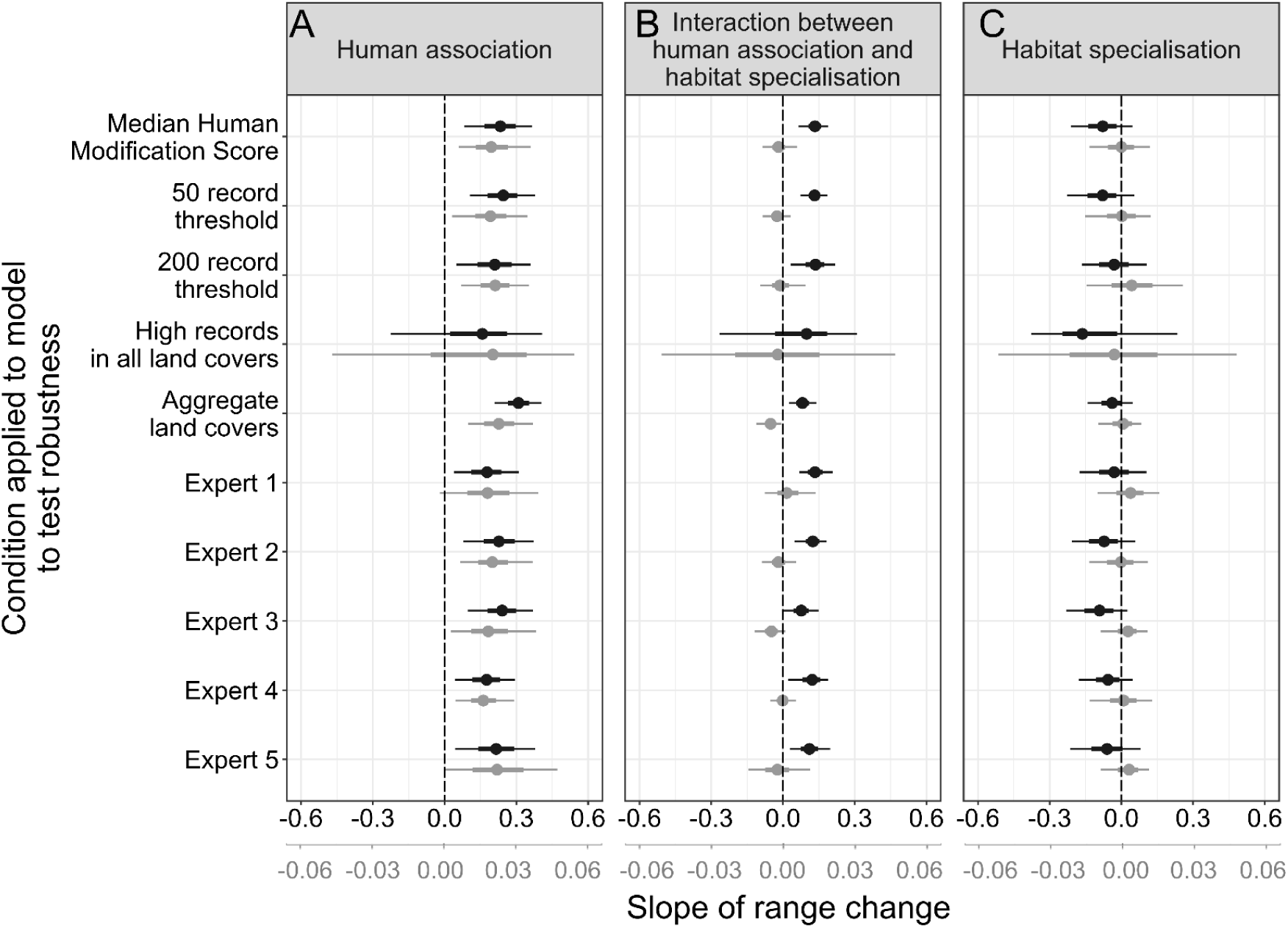
Effects of human association (A), habitat specialisation (C) and the interaction between human association and habitat specialisation (B) on range changes (slope values) by British invertebrate species between 1981-2000 and 2001-2020, comparing the effects of various tests for robustness. These include: These include: 1) Adjusting the thresholds for species selection to species with 50 presence records and 200 presence records (as compared to the 100 presence record threshold used in the main; top row median, here), 2) Adjusting the threshold for species selection to species with 15 records (presence or absence) in each of the 18 land cover classes, 3) Aggregating land cover classes according to the UKCEH Aggregate Land Cover Classes (Morton 2020) (Supp. Table 2), 4) Calculating human association index based solely on the human modification scores of each expert in turn (see Methods). Slope values are derived from a hierarchical Bayesian model with slope and intercept as a random effect by taxonomic group; applied to two measures of range change. Relative distribution change is in black, while change in absolute occupancy is in grey. Positive slopes in A would indicate that human associated species increased more than human avoidant species; negative slopes in C would show that habitat specialists declined most; and positive slopes in B that species specialising on the most human-modified habitats expanded most rapidly (only seen for the relative change metric). Error bars represent the 66% credible interval and the 95% credible interval. We observe very little influence of each treatment on the slopes of our model, giving us confidence in the robustness of our findings.

**Supplementary Figure 3:**
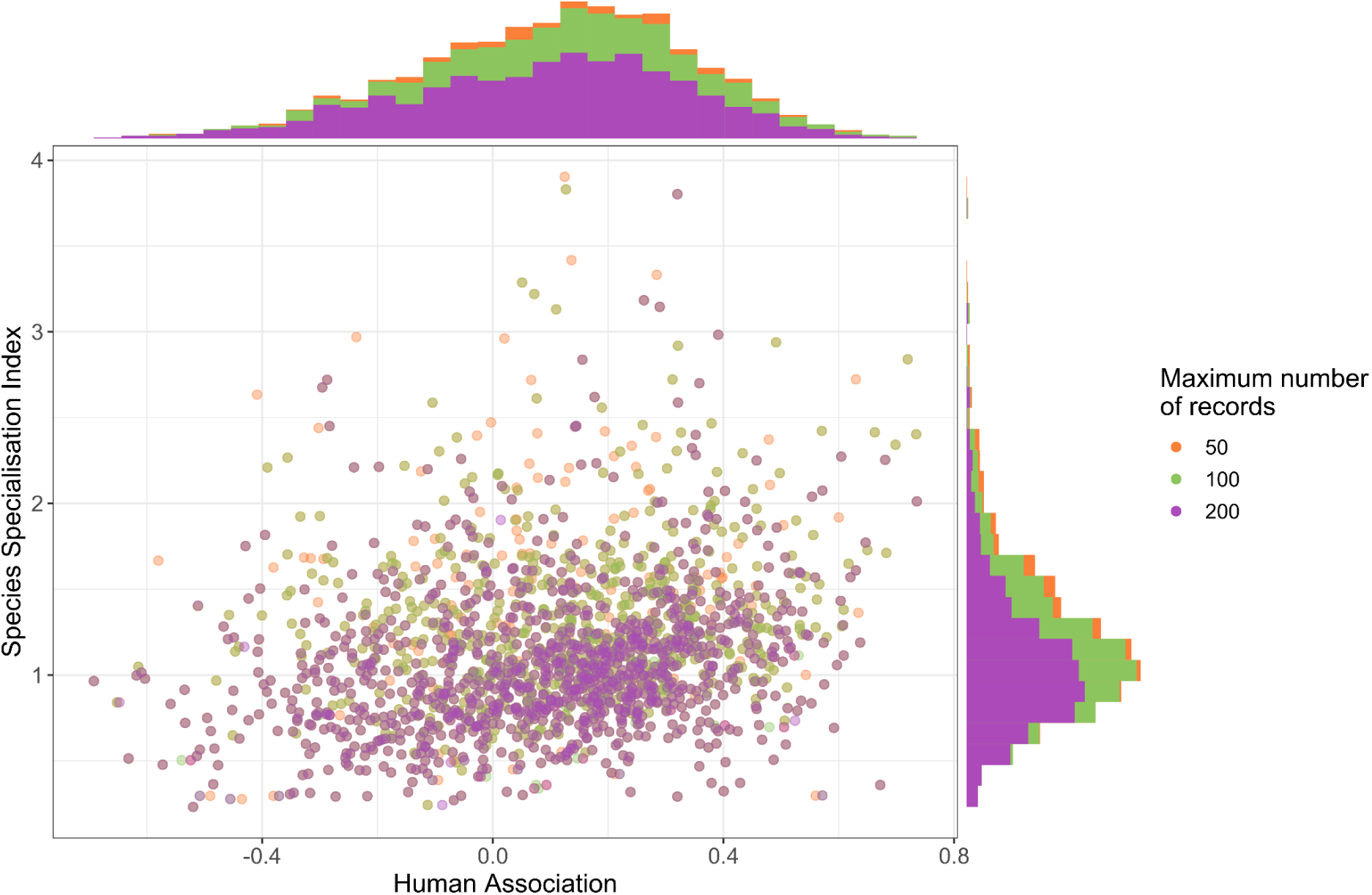
Distribution of species included by different recording filters of 50, 100 and 200 presence records per species. With a stricter recording focus (200 presence records), there is a bias towards species being more generalist, but no bias based on human association.

**Supplementary Figure 4:**
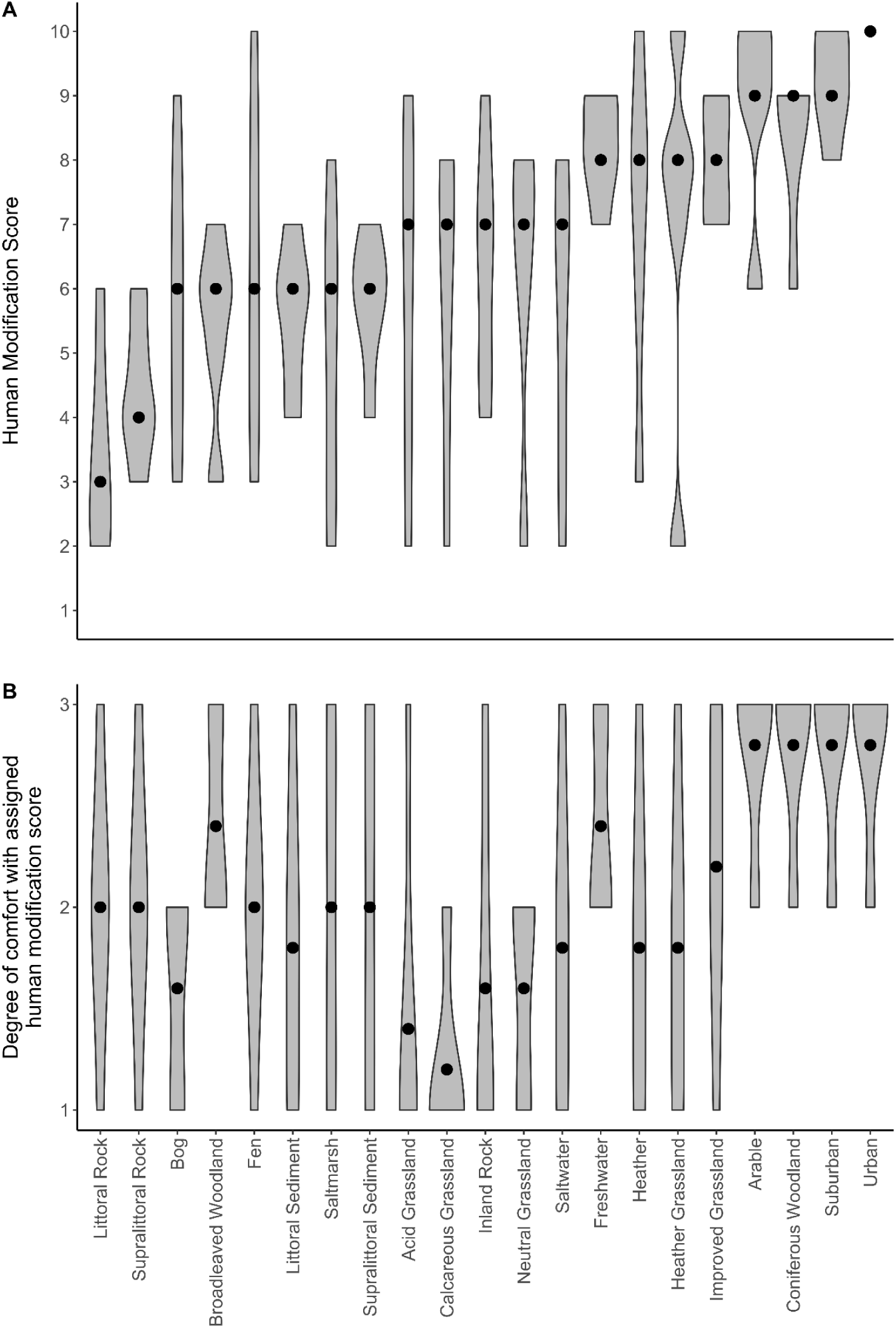
Human modification scores derived from expert elicitation interviews (A), in which 0 represents untouched by humans, and 10 represents completely changed by humans. Points represent the median scores of the five experts interviewed, which were used for further analysis. Interviewee comfort with their individual scores range from 1, being ‘very uncomfortable’, to 3, being ‘very comfortable’ (B). Points represent the median comfort for interviewees.

**Supplementary Figure 5:**
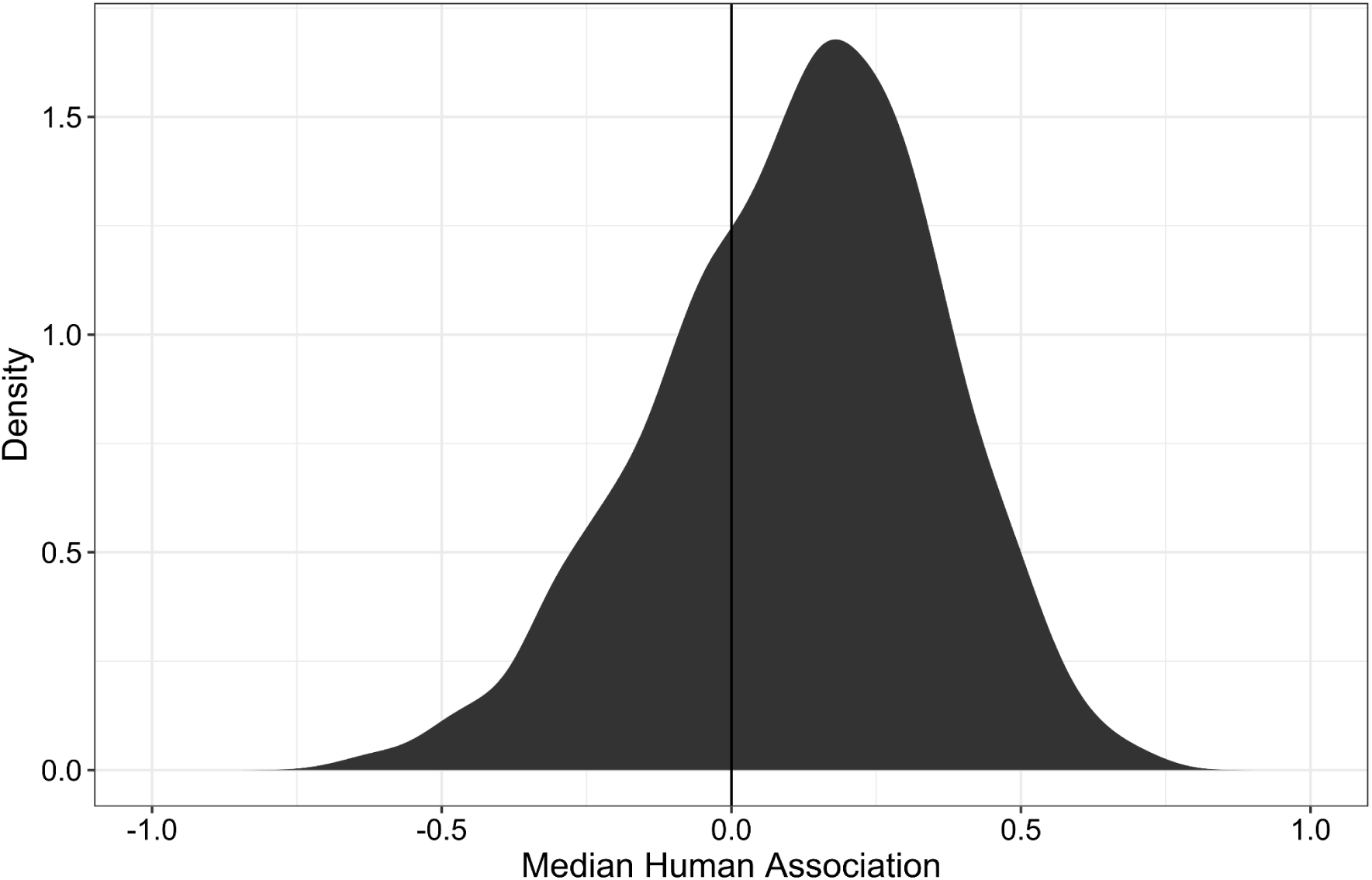
Density distribution of calculated human association scores, unscaled, from 1,722 species of invertebrates in Great Britain. There was a generally normal distribution, with 69% of species being identified as more associated with human modified land than not.

**Supplementary Figure 6:**
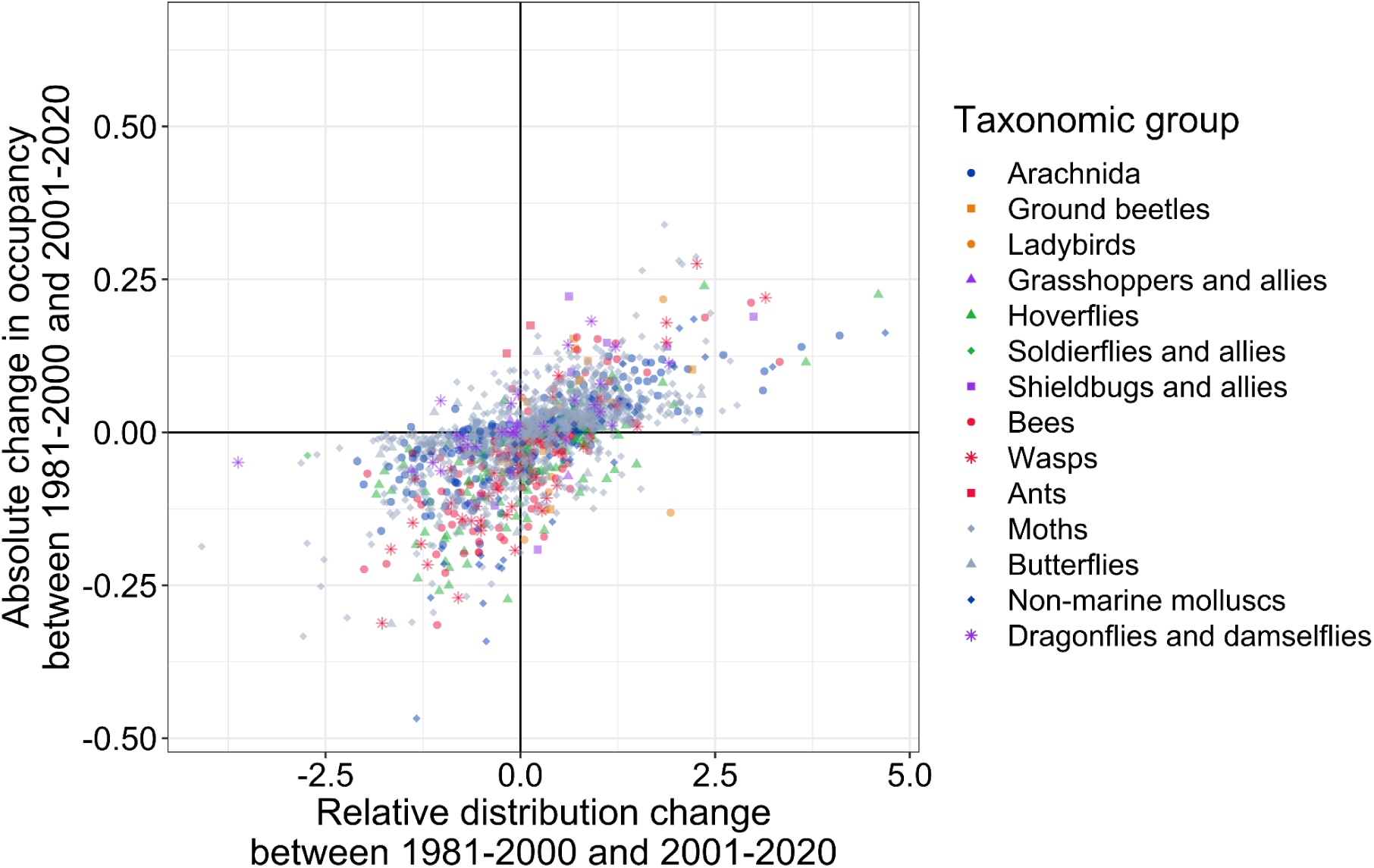
Relationship between absolute change in occupancy and relative distribution change, showing a mostly linear relationship, R=0.63, R^2^ = 0.4

**Supplementary Figure 7:**
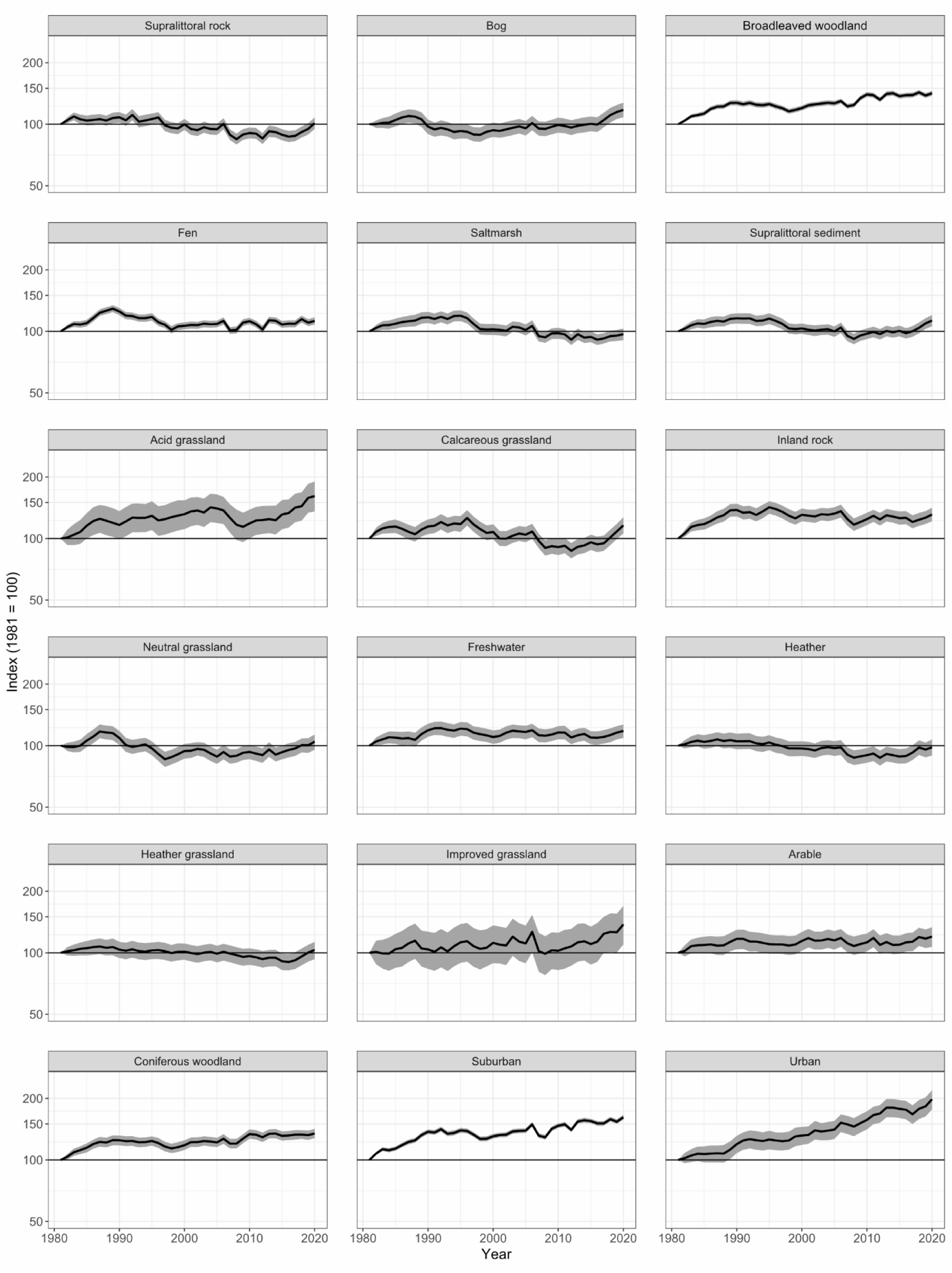
Arithmetic mean (line, 95% CI dark grey) of change in log odds occupancy for species, divided into land cover classes by the environment they are most and second most likely to be found in. Change in occupancy was calculated using the lambda indicator in the BRCIndicators package (Isaac et al. 2019; August et al. 2021). Numbers of species and change trends for each land cover type may be found in Supp. Table 3.

**Supplementary Figure 8:**
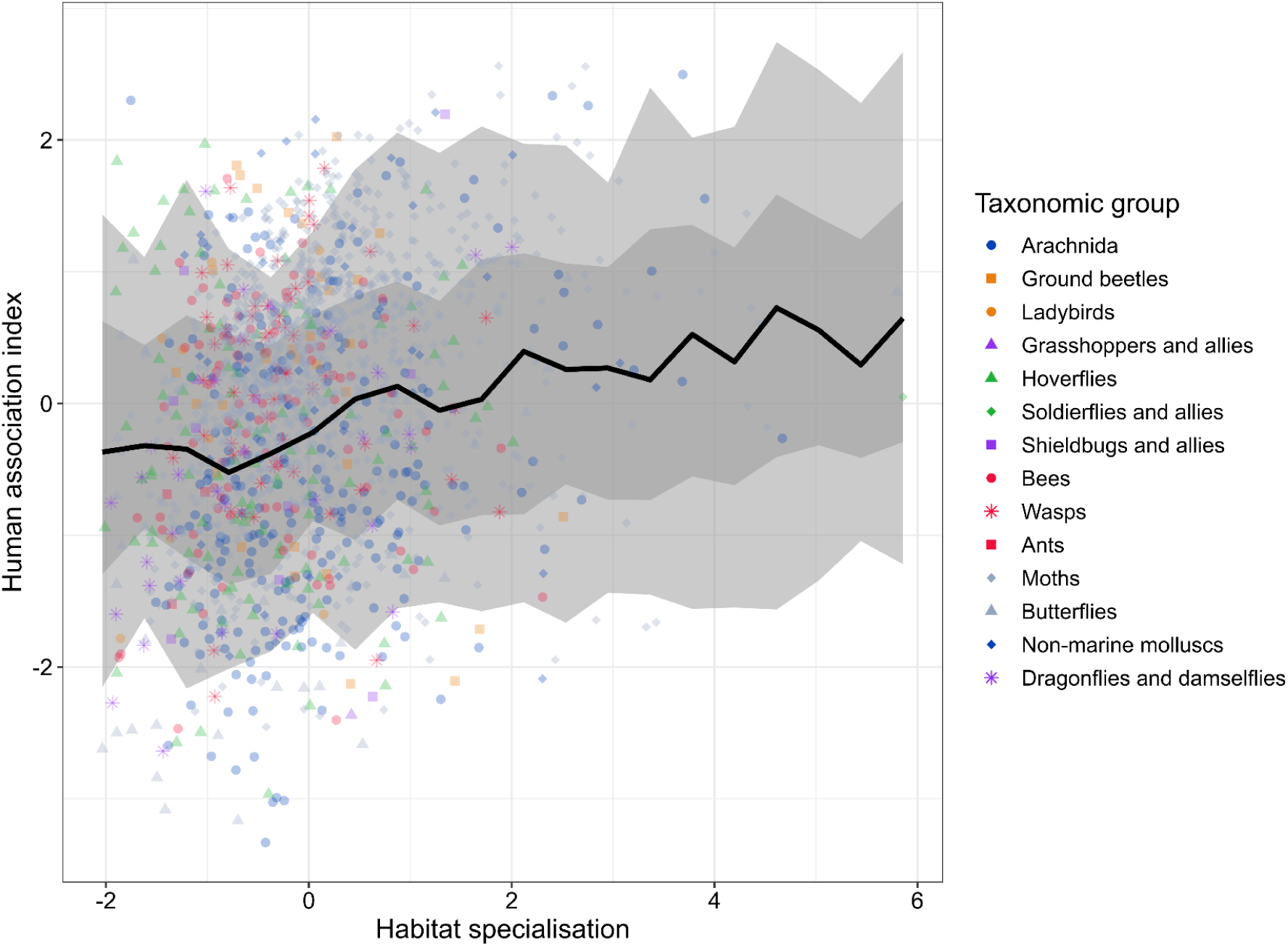
Point graph of human association index versus habitat specialisation, with slope predicted by Bayesian hierarchical modelling. Axes are scaled. The error ribbon represents the 66% credible interval and the 95% credible interval.

**Supplementary Figure 9:**
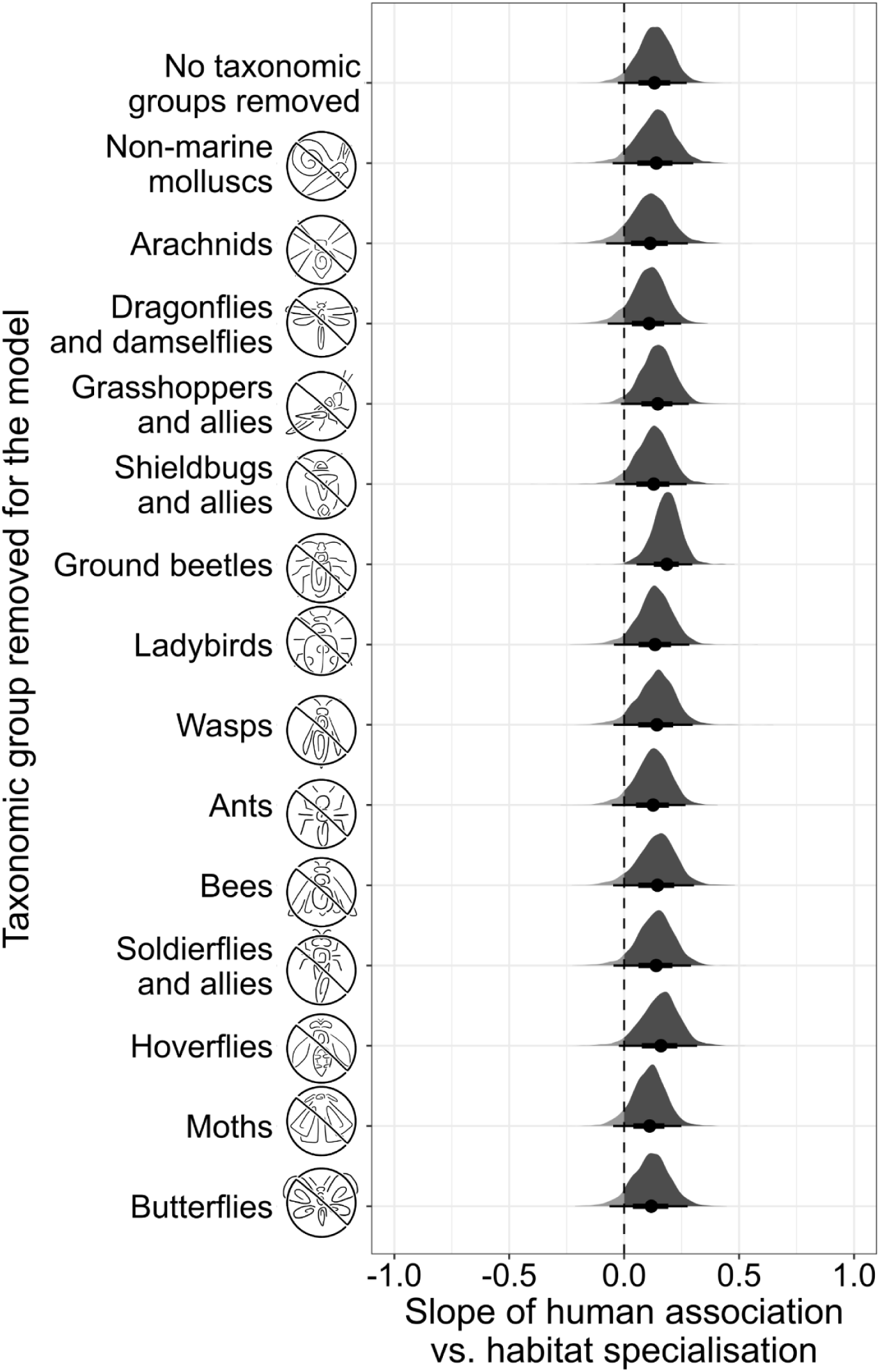
Change in the pooled slope of human association vs habitat specialisation when each taxonomic group is excluded in turn. There is very little difference between the results, suggesting that no one taxonomic group is driving the trends seen.

**Supplementary Figure 10:**
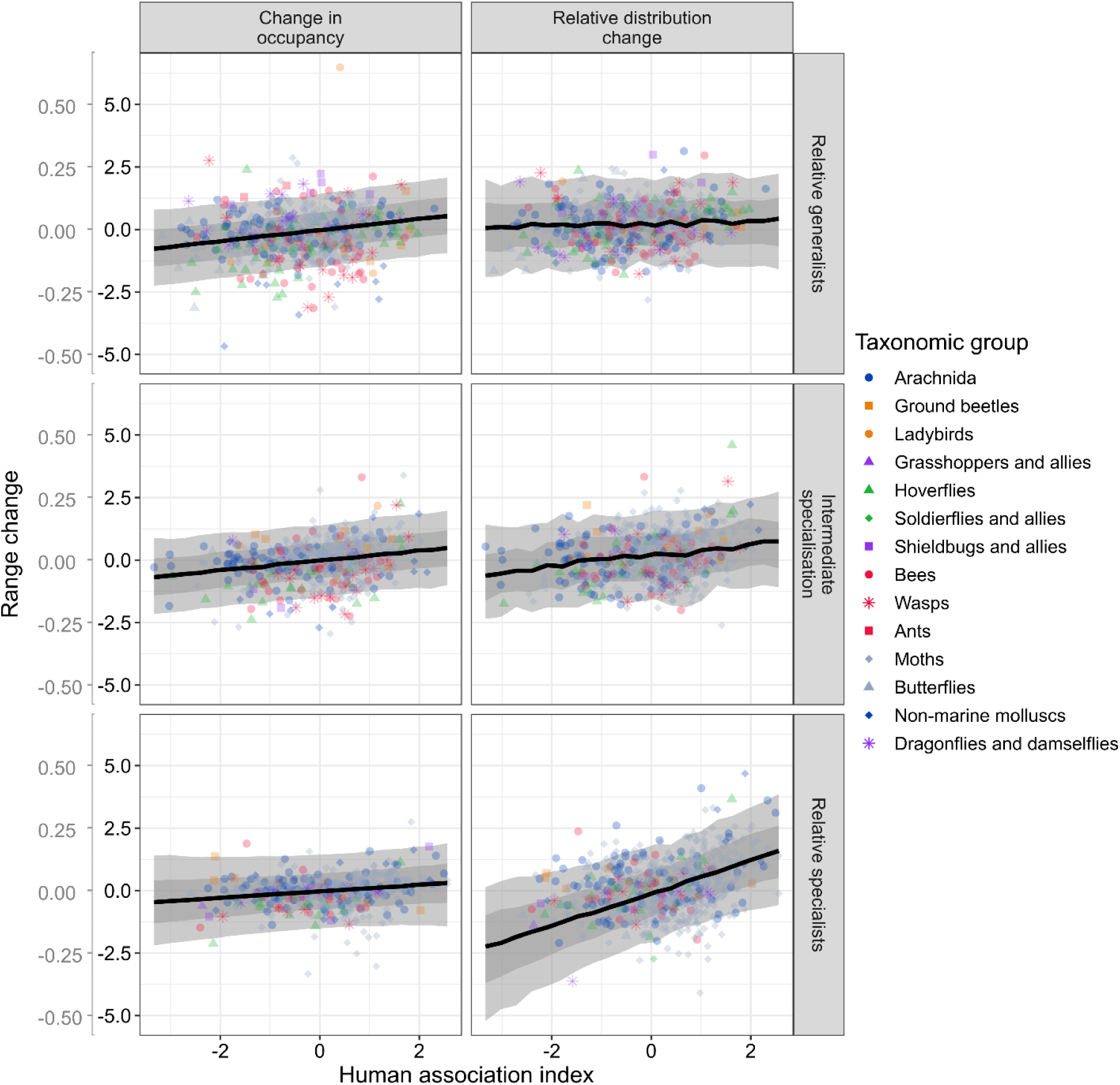
Point graphs of the relationships between range change (through relative distribution change, in black on the y axis, and absolute change in occupancy, in grey on the y axis) versus human association index, divided into the most generalist third of species, species of intermediate specialisation, and the most specialist third of species using cut_number (as in Figure 2.3), with slopes predicted from our Bayesian hierarchical model as used in Figure 2.2. The positive interaction between habitat specialisation and human association index for relative distribution change (B) shows an increasing slope as specialisation increases, but this is not also seen for absolute change in occupancy (A). The error ribbon represents the 66% credible interval and the 95% credible interval.

**Supplementary Figure 11:**
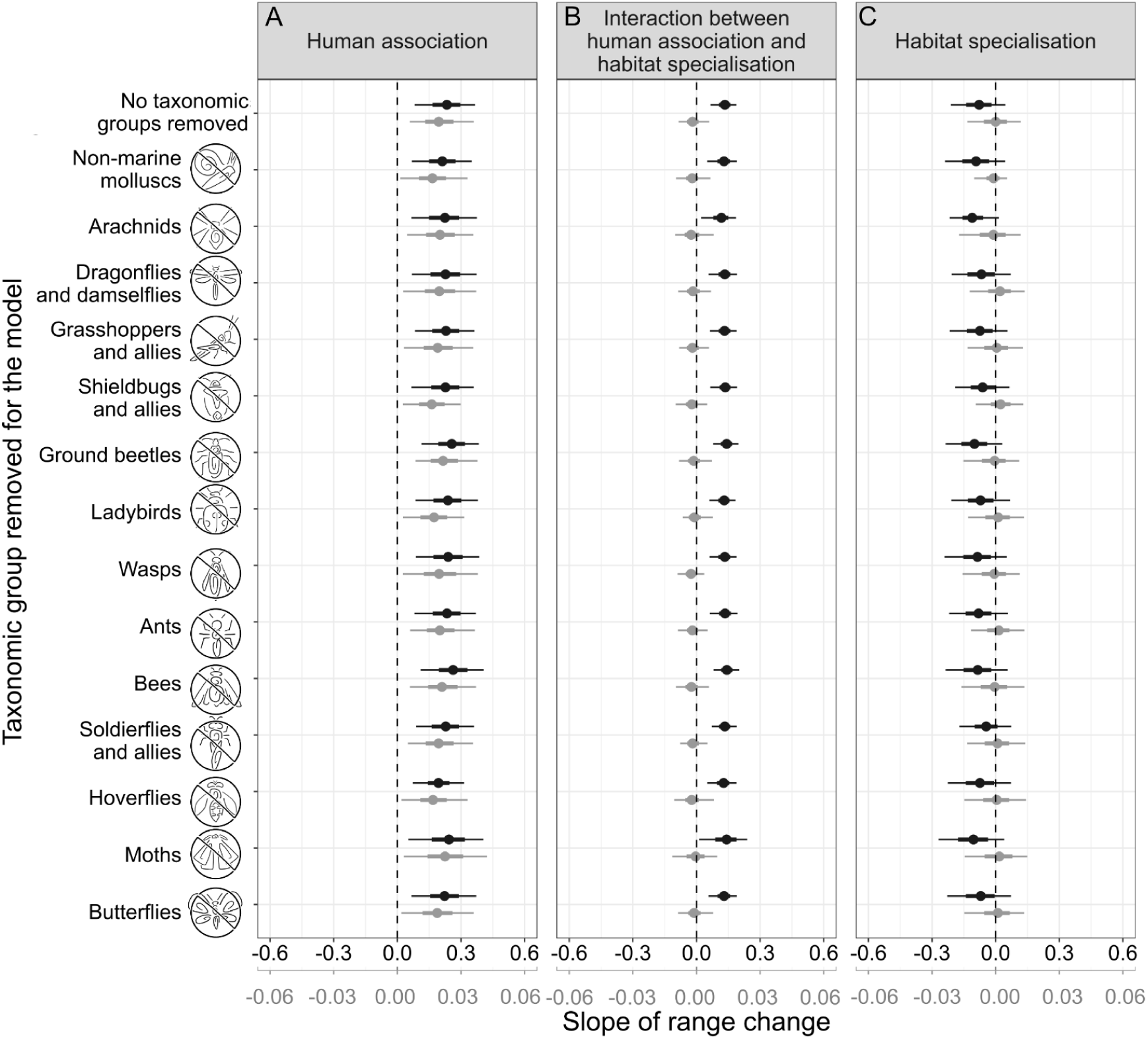
Difference between pooled slopes when each taxonomic group is excluded in turn, in terms of (A) the relationship between human association and range change, (C) the relationship between habitat specialisation and range change and (B) the interaction between the two. Relative distribution change is in black, while change in absolute occupancy is in grey. There is very little difference between the results, suggesting that no one taxonomic group is driving the trends seen. Slope values are derived from a hierarchical Bayesian model with slope and intercept as a random effect by taxonomic group; applied to two measures of range change. Relative distribution change is in black, while change in absolute occupancy is in grey. Positive slopes in A would indicate that human associated species increased more than human avoidant species; negative slopes in C would show that habitat specialists declined most; and positive slopes in B that species specialising on the most human-modified habitats expanded most rapidly (only seen for the relative change metric). Error bars represent the 66% credible interval and the 95% credible interval.

**Supplementary Figure 12:**
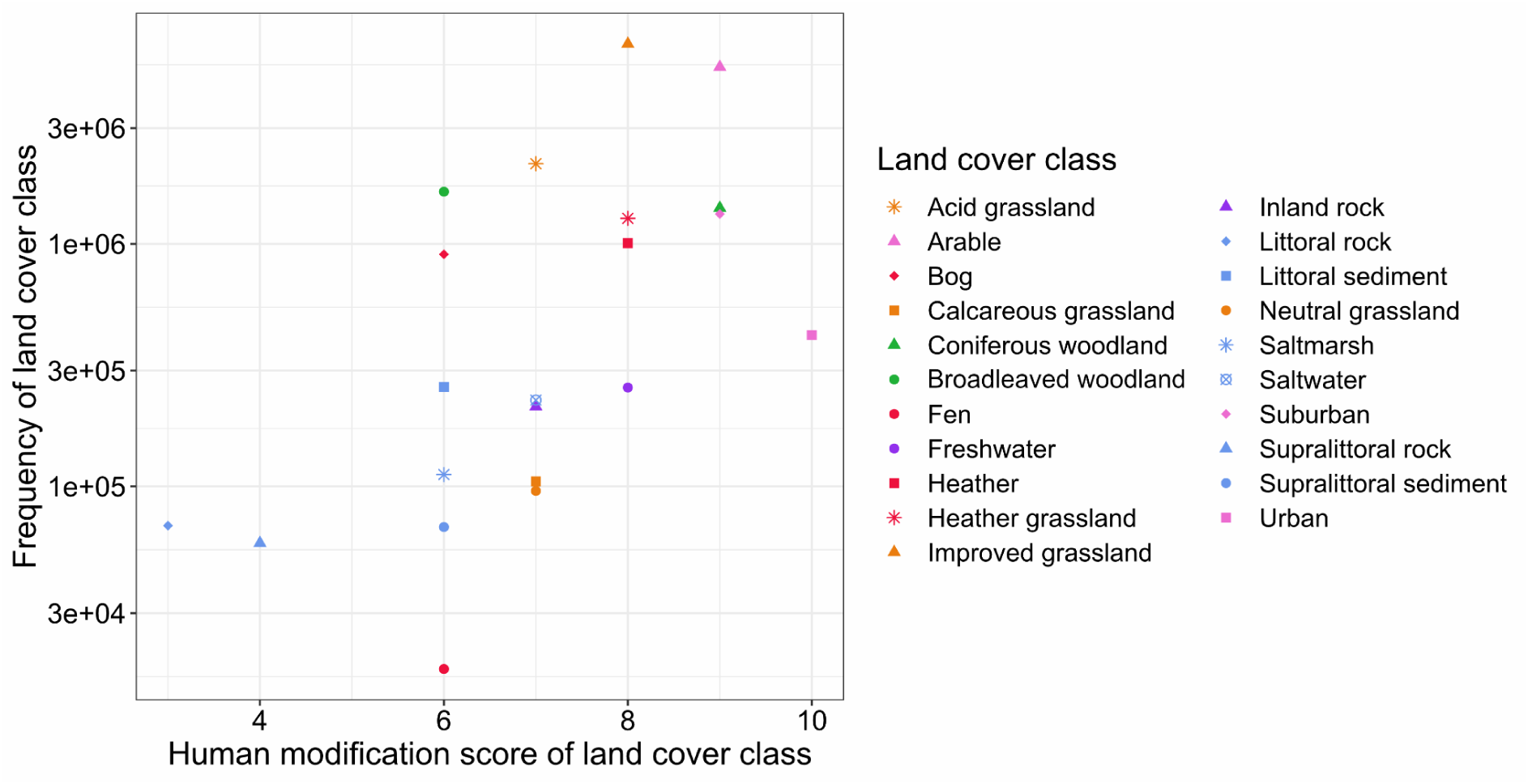
Relationship between human modification scores and the log number of hectares for each land cover class in Great Britain. R=0.59, R^2^ = 0.35

**Supplementary Figure 13:**
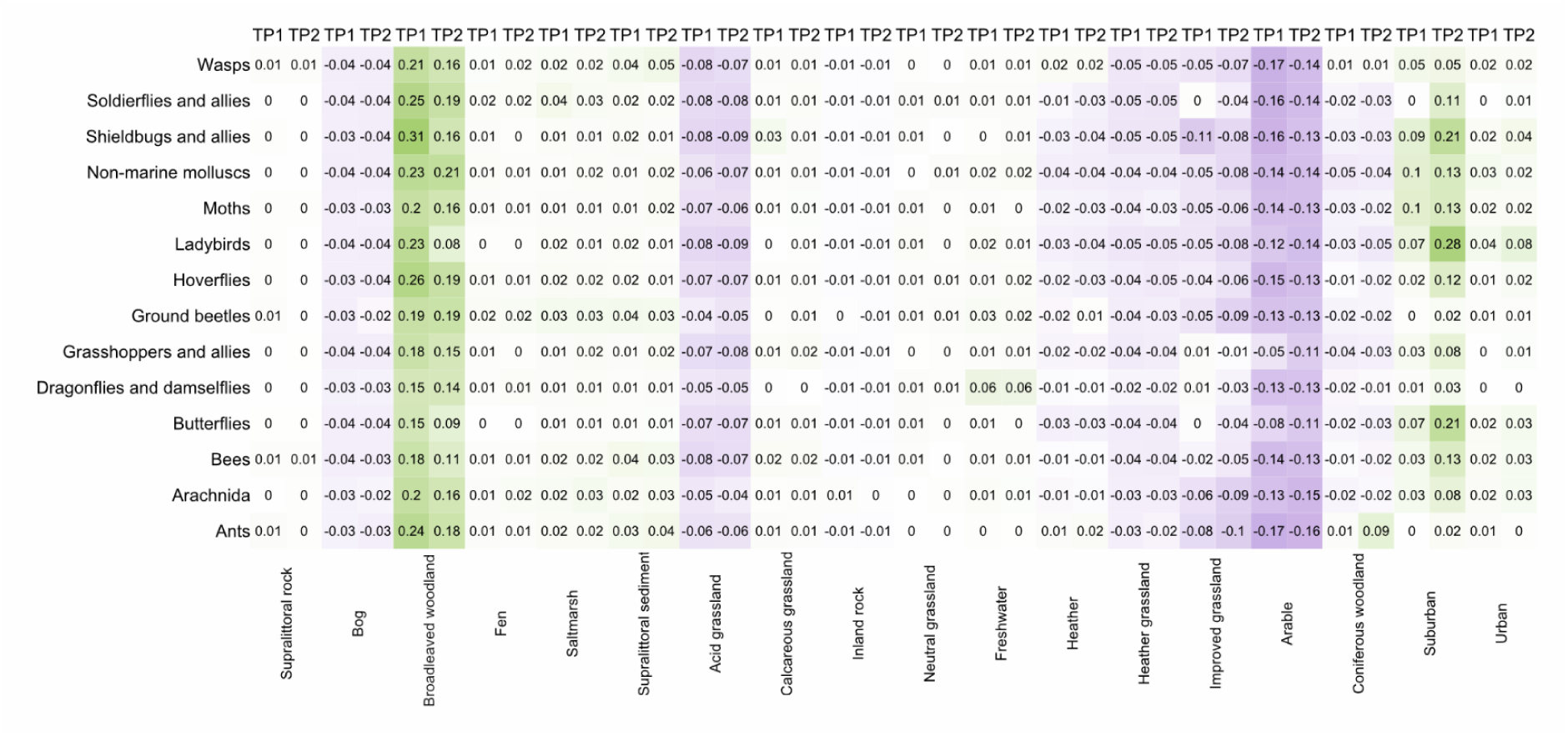
Levels of recording of different land cover types by each recording scheme for the two time periods. Levels of recording are relative to the overall availability of that land cover type in Britain (green indicates proportional over-representation; purple proportional under-representation). Each numerical value is the proportion of recordings for each land cover class (for a taxonomic group), minus the proportion of hectares for that land cover class across the whole of Great Britain (from the UKCEH land cover map; with littoral rock, littoral sediment and saltwater removed from the totals in both cases). TP1 = Time Period 1 (1981-2000), TP2 = Time Period 2 (2001-2020). Land covers are arranged from least modified (left) to most human-modified (right). Of the top eight most human-modified environments, three are over-recorded relative to land cover area (particularly suburban, and urban and freshwater to a lesser extent), while the remaining five are under-recorded. In contrast, of the eight least modified land covers, six are over-recorded relative to land area (especially broadleaved woodland) while two are under-recorded (bog, acid grassland).

**Supplementary Figure 14:**
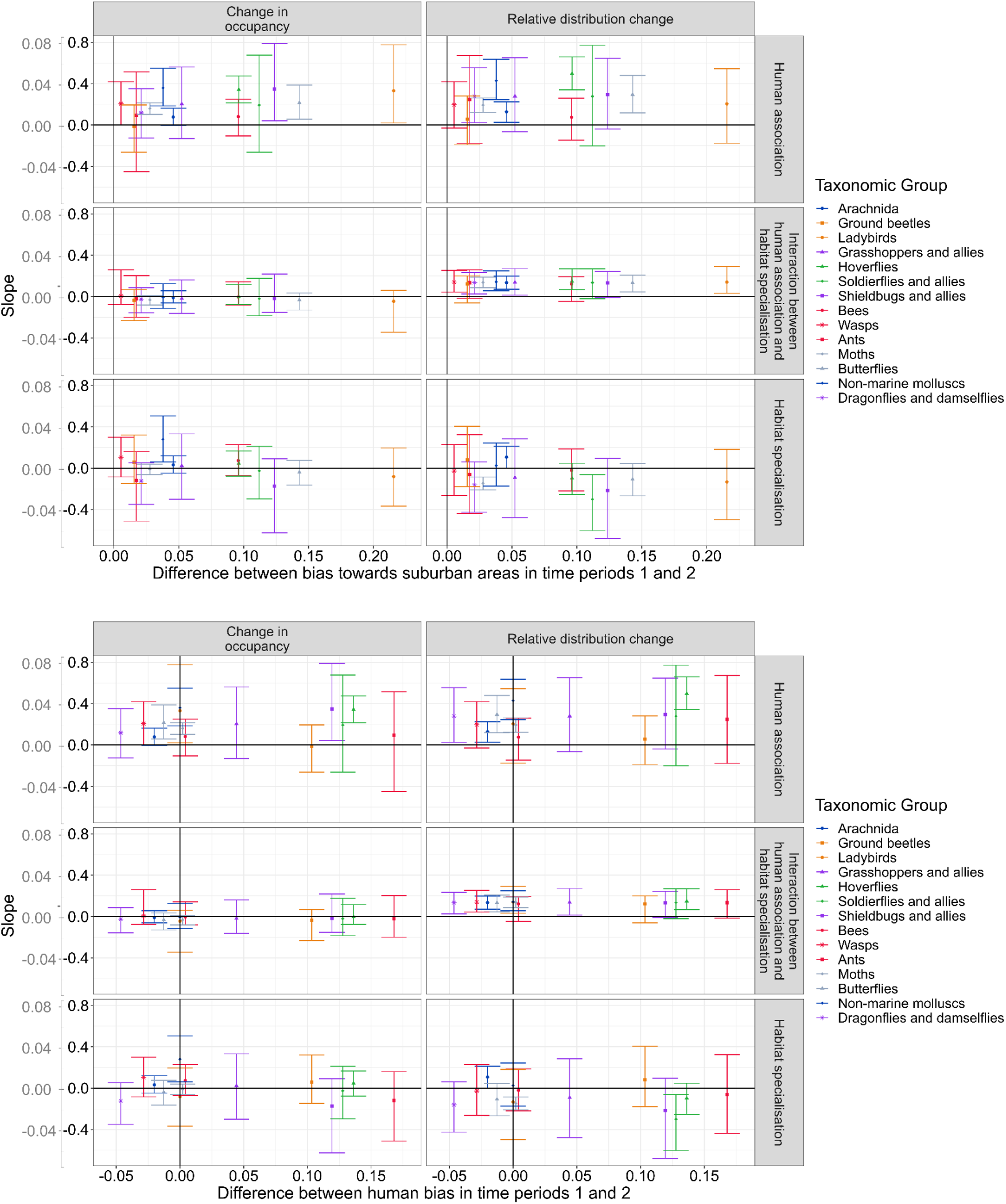
Relationship between the strength of the distribution change versus human association relationship (Slopes from Figure 2A, change in occupancy in grey and relative distribution change in black on the y axis) as a function of changes in recording between time periods 1 and 2 (Time Period 2 - Time Period 1 relative recording from Supp. Fig. 13). Upper panels show slopes in relation to increases in suburban recording. Lower panels show slopes in relation to changes in recording effort across all land cover types. Bias (x -axis) towards human environments was calculated as a Spearman’s correlation between human modification score for the land cover type, and divergence of records from the distribution of land cover types in the UKCEH land cover map. While there has been an increase in recording in human modified environments for many species, there is no correlation between the level of recording bias and the results. Each spot represents a different taxonomic group, with error bars representing the 95% CRI for these slopes.

**Supplementary Figure 15:**
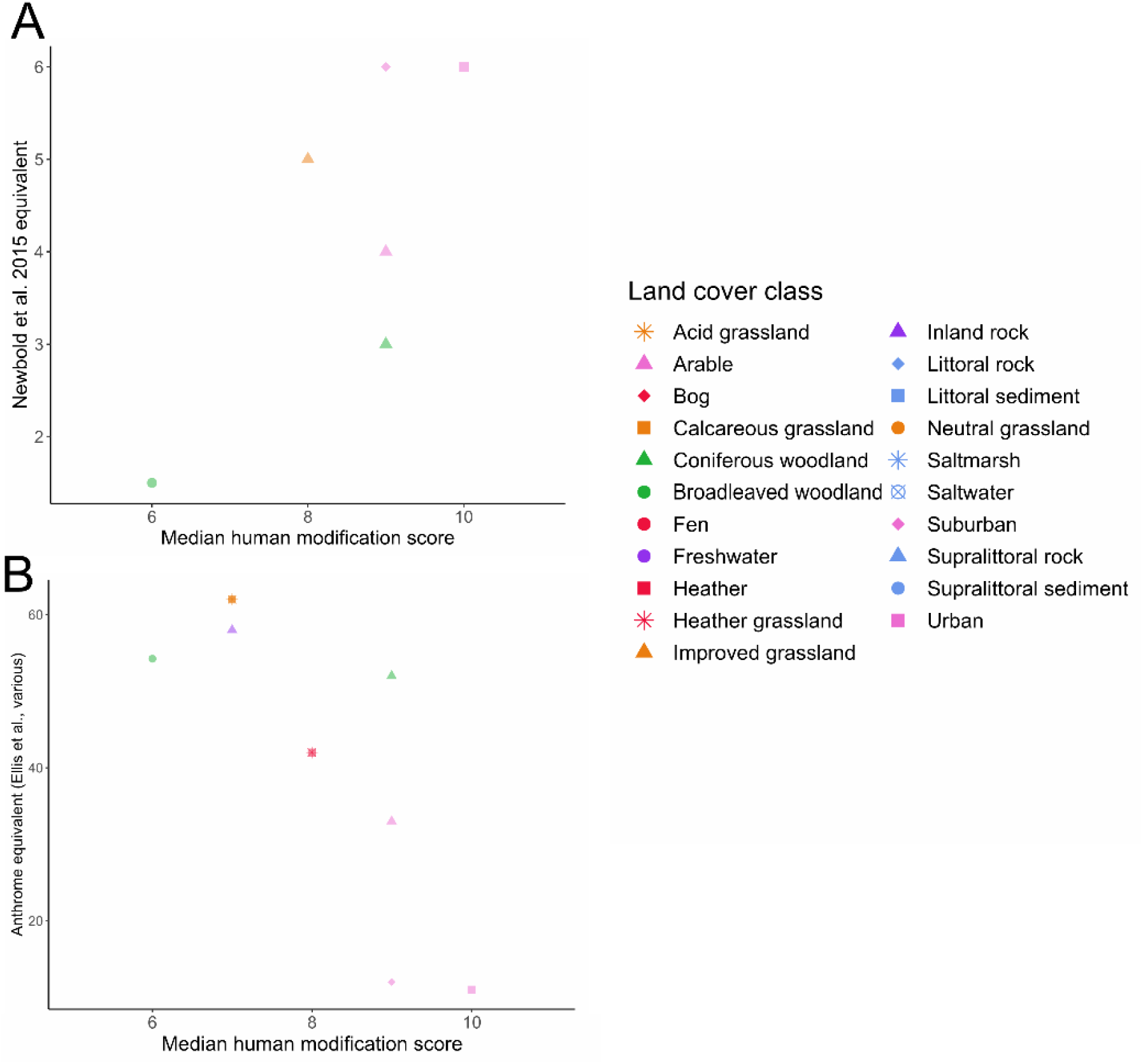
Comparisons of median human modification scores by land cover class (as in Supp. Fig. 4) to equivalent global rankings of human modification. A) refers to Newbold et al. (2015), and B) refers to anthromes, derived from several Ellis papers (Ellis and Ramankutty 2008; Ellis et al. 2020).

**Supplementary Table 2.1:** Steps in the methodology, showing the hectares and species removed at each step. ^a^ Species with no probability of presence for a land cover class were species with no pseudo-absence *or* presence records in that land cover class. ^b^ Species with no absolute occupancy values were species that lacked enough sites with records in at least two different years to be included in the occupancy modelling. ^c^ The species with no value for relative distribution change were *Harmonia axyridis*, *Leptoglossus occidentalis* and *Bombus hypnorum*, three non-native species with their first records being after 2000. Without records in the time period of 1981-2000, their values for relative distribution change could not be calculated.

| Step | Hectares lost | Remaining hectares | Species lost | Remaining species |
| --- | --- | --- | --- | --- |
| Total | 0 | 23621905 | 0 | 4698 |
| Remove hectares lacking any records | 22553531 | 1068374 | 0 | 4698 |
| Remove littoral habitats and saltwater | 17233 | 1051141 | 13 | 4685 |
| Remove hectares where the hectare contains two or more land cover types | 517754 | 533387 | 139 | 4546 |
| Remove hectares which experienced land use change | 133816 | 399571 | 76 | 4470 |
| Remove species with fewer than 100 presence sites |  |  | 2331 | 2139 |
| Remove species with no probability of presence for a land cover class<sup>a</sup> |  |  | 417 | 1722 |
| Species with no occupancy change value removed from total of 1722<sup>b</sup> |  |  | 98 | 1624 |
| Species with no value for relative distribution change removed from total of 1722<sup>c</sup> |  |  | 3 | 1719 |

**Supplementary Table 2.2:** UKCEH aggregate land cover classes used in this study (adapted from Morton et al. 2020)

| Aggregate Land Cover Class | Land Cover Class |
| --- | --- |
| Broadleaf woodland | Broadleaved woodland |
| Coniferous woodland | Coniferous woodland |
| Arable | Arable and horticulture |
| Improved grassland | Improved grassland |
| Semi-natural grassland | Neutral grassland |
|  | Calcareous grassland |
|  | Acid grassland |
|  | Fen |
| Mountain, heath and bog | Heather |
|  | Heather grassland |
|  | Bog |
|  | Inland rock |
| Freshwater | Freshwater |
| Coastal | Supralittoral rock |
|  | Supralittoral sediment |
|  | Saltmarsh |
| Built-up areas and gardens | Urban |
|  | Suburban |

**Supplementary Table 2.3:**
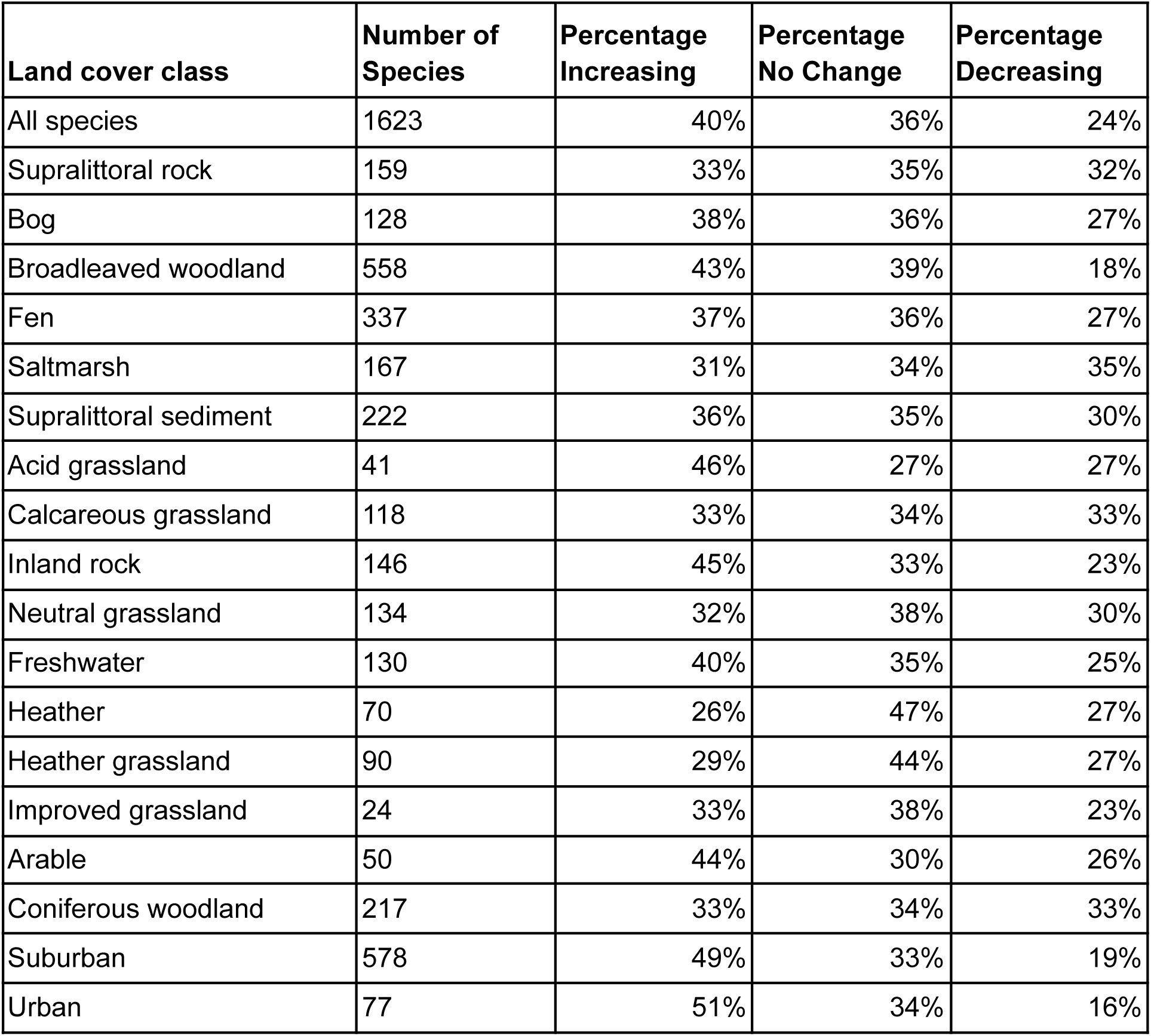
Summary statistics for Supp. Fig. 7. Numbers of species are those with the strongest and second strongest association with each land cover type. Rounded percentages of species experiencing increases, decreases and no change according to the lambda indicator are shown. ‘Decreases’ were species with a percent change per year of or more negative that −1.14, while ‘increases’ were species with a percent change per year of > 1.16 (derived from the species_assessment () function in the BRCIndicators package (Isaac et al. 2019; August et al. 2021)). The table also shows inter-specific variation in lambda indicators versus intra-specific variation in lambda indicators, for species associated with each land cover and for all species in total.

**Supplementary Table 4:** Effect Sizes of range change (r) vs. human association (h). As human association was scaled in the model, increases in human association are quoted as a multiple of the standard deviation (sd) of human association; that is, 0.24. Unscaled, human association ranges from −0.69 to 0.74. For *relative* distribution changes, comparing species, a 1 standard deviation (sd = 0.24) greater species’ human association index is associated with an increase in relative distribution change of 0.23 (on a scale of around −4 to 5). However, the median relative distribution change across all species is 0.17, meaning that the 0.23 increase per 1 standard deviation is 35% higher than this (Supp. Table 4). *Absolute* distribution change is easier to interpret. A 1 standard deviation (0.24) greater species’ human association index represented a 2% increase in the percentage of occupied grid cells relative to a less human-associated species.

| **Metric of r** | **Median Slope**<br>**(change in r**<br>**per 1 sd h)** | **Median r at h =**<br>**-2sd** | **Median r**<br>**(across all**<br>**species)** | **Median r at h =**<br>**2sd** |
| --- | --- | --- | --- | --- |
| Relative<br>distribution change | 0.23 | -0.29 | 0.17 | 0.63 |
| Absolute change in<br>Occupancy | 0.02 | -0.041 | -0.001 | 0.039 |

Supplementary Tables 5 - 50: Model coefficients for all model iterations

